# Neural populations in the dorsal and ventral cervical spinal cord share temporal dynamics during reach

**DOI:** 10.64898/2026.10.12.758656

**Authors:** Trevor S. Smith, Shiro Egawa, Akito Kosugi, Kazutaka Maeda, Chiaki Sugai, Amit Yaron, Kazuhiko Seki

## Abstract

Neural population dynamics provide a popular framework for studying motor cortex function. However, whether such dynamics reflect stage-specific computations or are shared across the input and output stages of a sensorimotor circuit is unknown. In the spinal cord, afferent inputs and motor outputs are segregated by anatomical depth. Here we developed depth-resolved Neuropixels recordings from the rat cervical spinal cord during targeted reaching behavior. Across our recordings, population activity was low-dimensional and organized into transient, rotational and target-specific dynamical motifs. Despite differences in single-neuron activity by depth, dorsal and ventral populations expressed nearly identical dynamics across targets and animals. The dominant target-invariant component was strikingly synchronized across depths, whereas other components, including the target-specific component, showed condition-dependent ventral-to-dorsal delays with timing consistent with movement-related feedback. Thus, cervical spinal population dynamics span sensory and motor territories, and may reflect shared coordination of activity rather than purely state-specific local processing.

## Introduction

Neural population activity often evolves through low-dimensional trajectories that have been interpreted as signatures of computation through dynamics^1^, particularly in the motor cortex. Such population-level structure, including rotational and transient motifs^2,3^, has been linked to movement generation and control. However, whether these dynamics reflect intrinsic circuit computation or are primarily driven by structured input and feedback remains an active area of debate.

While population dynamics have been extensively characterized in cortical circuits, far less is known about whether similar principles apply to spinal sensorimotor circuits. While this extension is being actively explored (e.g.,^4–7^), this gap is particularly relevant given the anatomical and functional organization of the spinal cord, where sensory-dominated dorsal regions and motor-dominated ventral regions are traditionally considered as serving distinct roles. If population dynamics reflect local computation, one might expect these regions to exhibit different dynamical structures. Conversely, if such dynamics arise from shared inputs or network-level interactions, similar population activity may be observed across depth.

Directly addressing this question has been limited by technical constraints. Simultaneous, depth-resolved recording in the spinal cord during voluntary behavior has been difficult to achieve at sufficient scale and resolution to characterize neural population dynamics. As a result, it remains unclear whether spinal population activity exhibits low-dimensional structure comparable to cortex during skilled behavior, and whether such dynamics differ across anatomical partitions.

First, we introduce an approach for high-density, depth-resolved recordings from the cervical spinal cord of awake, behaving rats using Neuropixels^8^ probes. This method enables simultaneous sampling of large neural populations across dorsal and ventral regions during a skilled forelimb reaching task.

Next, we use principal component analysis (PCA), joint PCA^2^ (jPCA), and demixed PCA^9^ (dPCA) to address complementary aspects of population activity: PCA establishes the dimensionality of the neural activity, jPCA isolates structured temporal dynamics within that low-dimensional space, and dPCA separates shared temporal structure from condition-dependent modulation. Using these analyses, we identify reproducible low-dimensional population dynamics in the spinal cord, with highly similar temporal structure between dorsal and ventral partitions.

## Results

### An animal preparation for recording from the cervical spinal cord during behavior

Animals were implanted with a custom head plate (**Fig. 1A**) and spinal recording chamber (**Fig. 1B-F**) and were taught to reach with their right hand (**Fig. 1G**), to a lever placed either ipsilateral or contralateral to the reaching hand (**Fig. 1H**). After training and implants, neural data was recorded from the awake, behaving, animal in a single terminal experiment (**Fig. 1I**).

**Figure 1:**
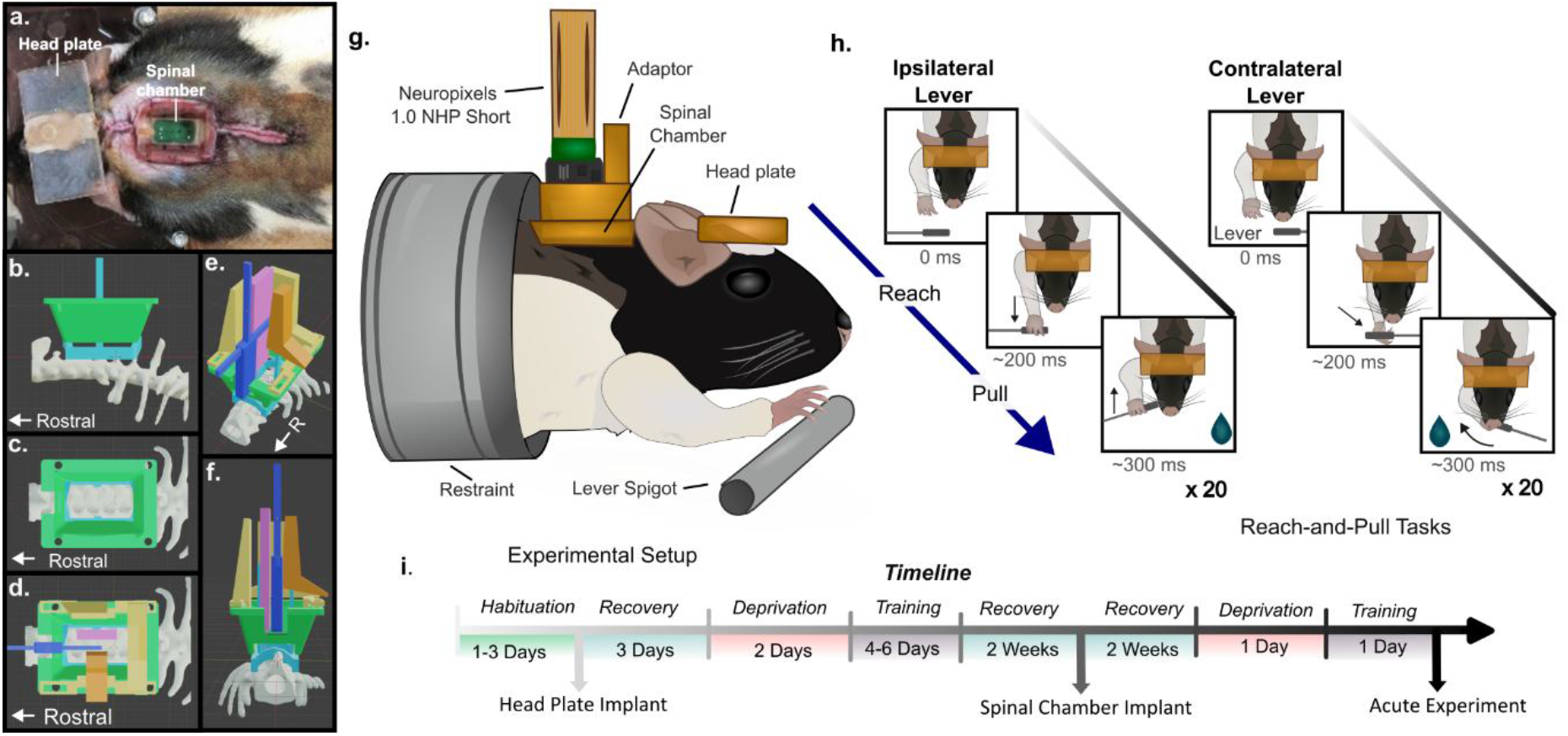
Experimental and recording paradigm. A) Experimental animals were fitted with both a head plate and spinal chamber for recording. **B-F**) A spinal chamber is implanted over the cervical vertebrae (C2-T2) to securely fasten the Neuropixels probe (Neuropixels 1.0 NHP Short) during recording. The spinal recording chamber and Neuropixels electrode adapter utilized a custom 3D-printed resin interface, anchored to the 2^nd^ cervical and 2^nd^ thoracic vertebrae. **G**) Diagram of recording paradigm. During the acute procedure, neural recordings are made from the cervical spinal cord while the animal performs reaches towards a lever. The head plate and spinal chamber interface with experimental apparatus to maintain recording stability. **H**) The animal is trained to reach to a lever placed either ipsilateral or contralateral to the reaching arm. **I**) The experimental timeline for hardware implants and training.

### Single Unit Analysis

After collecting neural data within our two-target reaching paradigm, further analysis was performed offline. Despite recording from the cervical spinal cord during awake behavior, momentary electrode drift was relatively minor, confined to ±50 µm, corresponding to 2 electrodes either direction on the Neuropixels 1.0 NHP Short probe (20 µm intercontact distance) within epochs of reaching trials (**Figure 2A**). To mitigate this small electrode drift, raw voltage traces were corrected with DREDge^10^ prior to spike sorting. Next, we used the depth-dependent local-field potentials of the spinal cord evoked by peripheral nerve stimulation to coarsely identify electrode channel depth (**Figure 2B-D**). Following drift correction, spike sorting, and unit curation (see **Sup. Fig. 1, Sup. Table 1**), we retained a total of 797 neurons (e.g., **Figure 2E**). Neural spike trains were then parsed into trials, aligned relative to hand takeoff from home position (**Figure 2F-G)**. Across our 8 rats, mean reach duration was 212 ± 109 ms to the ipsilateral and 249 ± 61 ms to the contralateral lever target. Individual rats exhibited similar kinematics trajectories across trials (**Figure 2H**). To capture this reaching epoch, we used an interval 500 ms before to 1000 ms after reaching onset for our peristimulus time histograms (PSTH), discretized with a 1 ms bin width. We then omitted neurons (n=154) which only fired outside this binned interval. Thus, a total of 643 neurons showed spiking activity during reaching trials and were retained for analysis.

**Figure 2:**
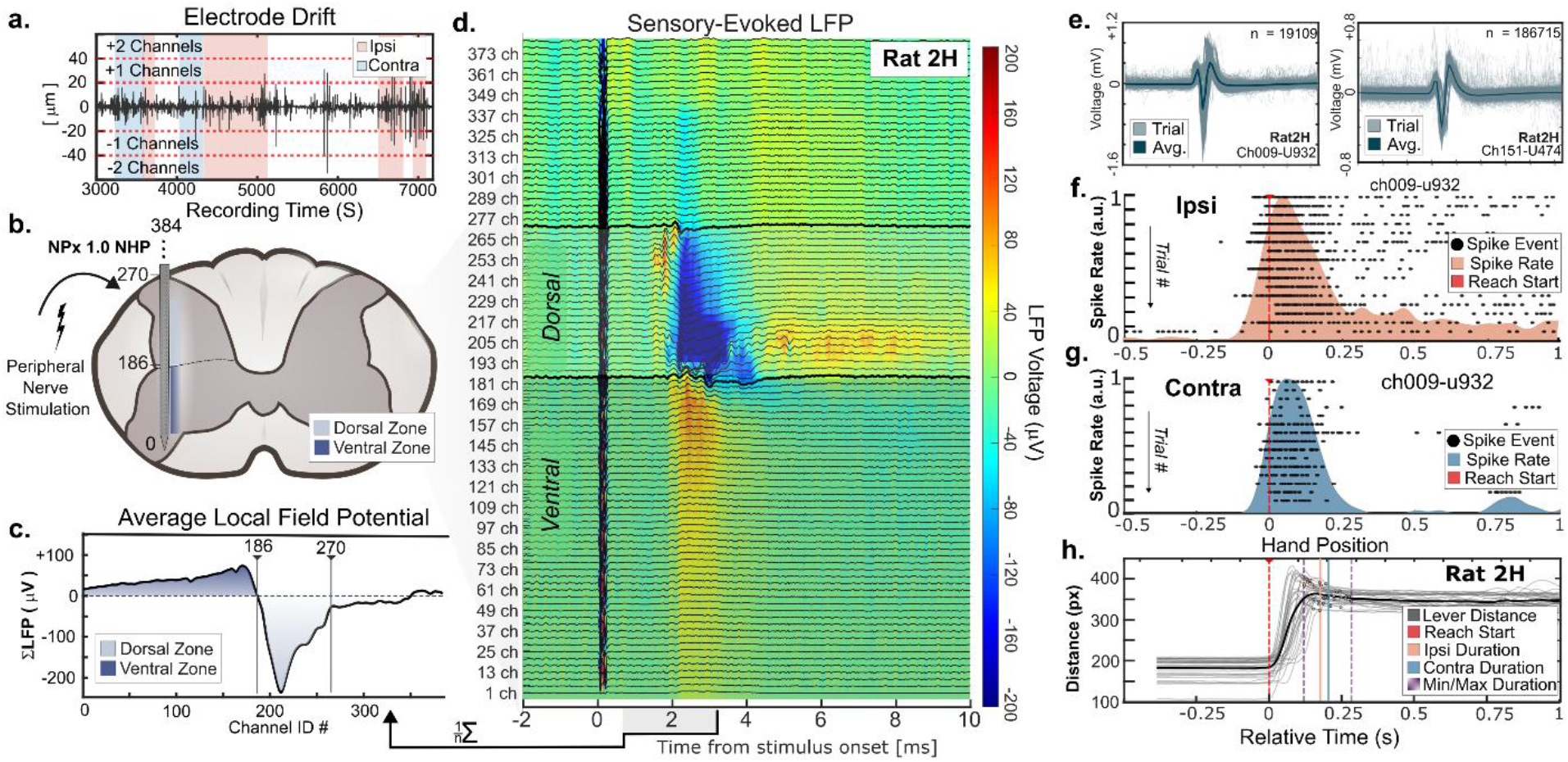
A) Electrode drift recorded during behavioral epochs (red = ipsilateral reach, blue = contralateral reach) (Here, reaching trials were performed within the ∼3000-7000 second period of the larger 2-hour recording). B) The spatiotemporal signature elicited by peripheral nerve stimulation may be used to assess channel depth. C) We assigned the dorsal-ventral border as the channel where the sign-change occurred relative to the LFP traces recovered in the 1-3 ms period after stimulation. D) The depth-dependent response of the electrode LFP displayed a spatial-temporal signature indicative of electrode placement. E, F) Neural spikes were richly extracted from our entire recorded paradigm. G) Neural responses in the C6 spinal cord were often tuned to motor responses (H). Reaching duration and kinematics were highly similar across trials within an animal.

**Table 1:** Canonical correlations between jPCA subspaces measured between the ensemble jPCA and the depth-dependent jPCA subsets. Significance was determined relative to a sample-size balanced set of 5000 shuffled surrogate statistics, randomly drawn across depth from the ensemble. Both dorsal and ventral subsets of neurons generated jPCA subspaces which were near-linear (ρ>0.98), however, ventral-only neuron jPCA structure was significantly more similar to the ensemble than the shuffle (i.e., ventral alone neurons were a better fit than an admixture of ventral and dorsal neural IDs).

| Grouping | Plane | Test Val<br>($\rho$) | Shuff<br>CI<br>(5%) | Shuff<br>CI<br>(95%) | pVal |
| --- | --- | --- | --- | --- | --- |
| Ventral | JP1 | 0.999 | 0.665 | 0.993 | **0.0004** |
|  | JP2 | 0.999 | 0.946 | 0.998 | **0.004** |
|  | JP3 | 1.000 | 0.976 | 0.999 | **0.001** |
| Dorsal | JP1 | 0.938 | 0.594 | 0.993 | 0.674 |
|  | JP2 | 0.997 | 0.915 | 0.998 | 0.074 |
|  | JP3 | 0.985 | 0.954 | 0.999 | 0.494 |

Spike rates were derived from the PSTHs by convolution with a 20 ms Gaussian smoothing kernel (**Figure 3A**). Per-unit ipsilateral-contralateral peak-peak timing plots of neural spike rates indicated that peak timing was loosely conserved for each neuron across motor conditions (**Figure 3B-C**). The distribution of peak activity nonetheless differed by motor condition (Kolmogorov-Smirnov; p=0.0086, D=0.0971, n=545596), with neuron peak activity firing later for the contralateral reaching condition across the ensemble (**Figure 3D**). Neuron-wise PSTH vectors were weakly correlated across the ensemble for ipsilateral and contralateral reaches (R: 0.247) (**Figure 3E**). Small differences in average excitability between ipsilateral and contralateral reach condition, evaluated as the root-mean square (RMS) of the spike rate, were observed in the 643-neuron ensemble (**Figure 3F**) (Rank-Sum; W=394617.5, N=643; p=0.004), however this difference disappeared when excluding the ∼23% (145/643) neurons (**Sup. Table 2**) which fired in only a single motor condition (Rank-sum; W= 244150, N=498; p=0.366). This suggests that while some neurons were active only in a single motor condition, neurons active in both conditions showed no population-wide difference in firing-rate magnitude between conditions.

**Figure 3:**
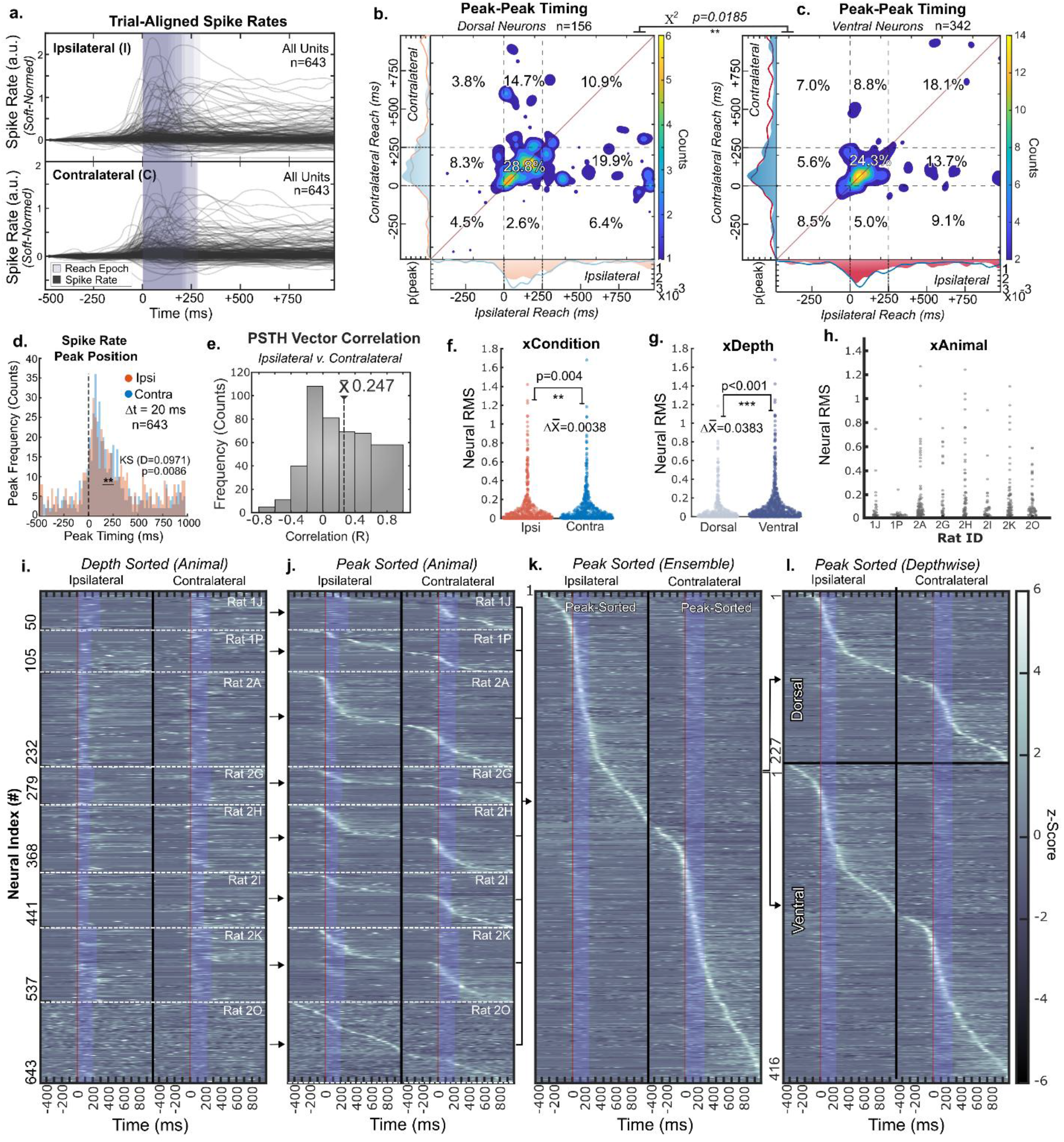
Neural tuning within and across rats. **A**) Average spike rates were collected for a total of 643 trial-active neurons across 8 rats. Rates were collected for reaches to both the ipsilateral or contralateral target. The average reaching epoch for each rat is overlaid in purple (Grand Mean: 212 ms ipsilateral; 249 ms contralateral). **B-C**) We considered the timing of peak spike rate independently for dorsal (n=156) and ventral (n=342) neurons which fired for both motor conditions. We considered a premotor period -500 to 0 ms before motor onset, the reaching period from 0 to +250 ms after motor onset, and a post-motor period from +250 to +1000 ms after motor onset. We evaluated the consistency of this peak timing for both conditions. Most neurons which maximally fired during the reaching epoch towards the ipsilateral lever also maximally fired during the reach towards the contralateral lever. The distribution of paired ipsilateral-contralateral peak timing differed by depth (χ2, p=0.012). **D**) The distributions of all neural peaks were not identical across reaching target (K-S test, p=0.0086, D=0.0971, n1=545, n2=596), but were coarsely similar. **E**) PSTH vectors (i.e., average firing rates) were weakly correlated between ipsilateral and contralateral reaches (R^2^=0.247). **F**) A small effect difference in ipsilateral and contralateral neural activity, measured as the root mean square (RMS) of the PSTH, was observed between ipsilateral and contralateral reaches. **G**) Ventral neurons had a higher activity level than dorsal neurons. **H**) Some variability in activity level was observed across animals. **I**) Baseline-subtracted, variance-normalized, depth-ordered spike rates across 8 rats. The rat-wise average reaching epoch is overlaid in purple. **J**) Per-rat, peak-sorted spike rates showed a smooth continuum of peak neural timing across the ensemble, with enrichment during the motor period (+0-250 ms). **K**) Ensemble (pooled) peak-sorted spike rates from 643 neurons across all rats. **L**) Peak-sorted spike rates of ensemble data, grouped by the depth profile.

**Table 2:**
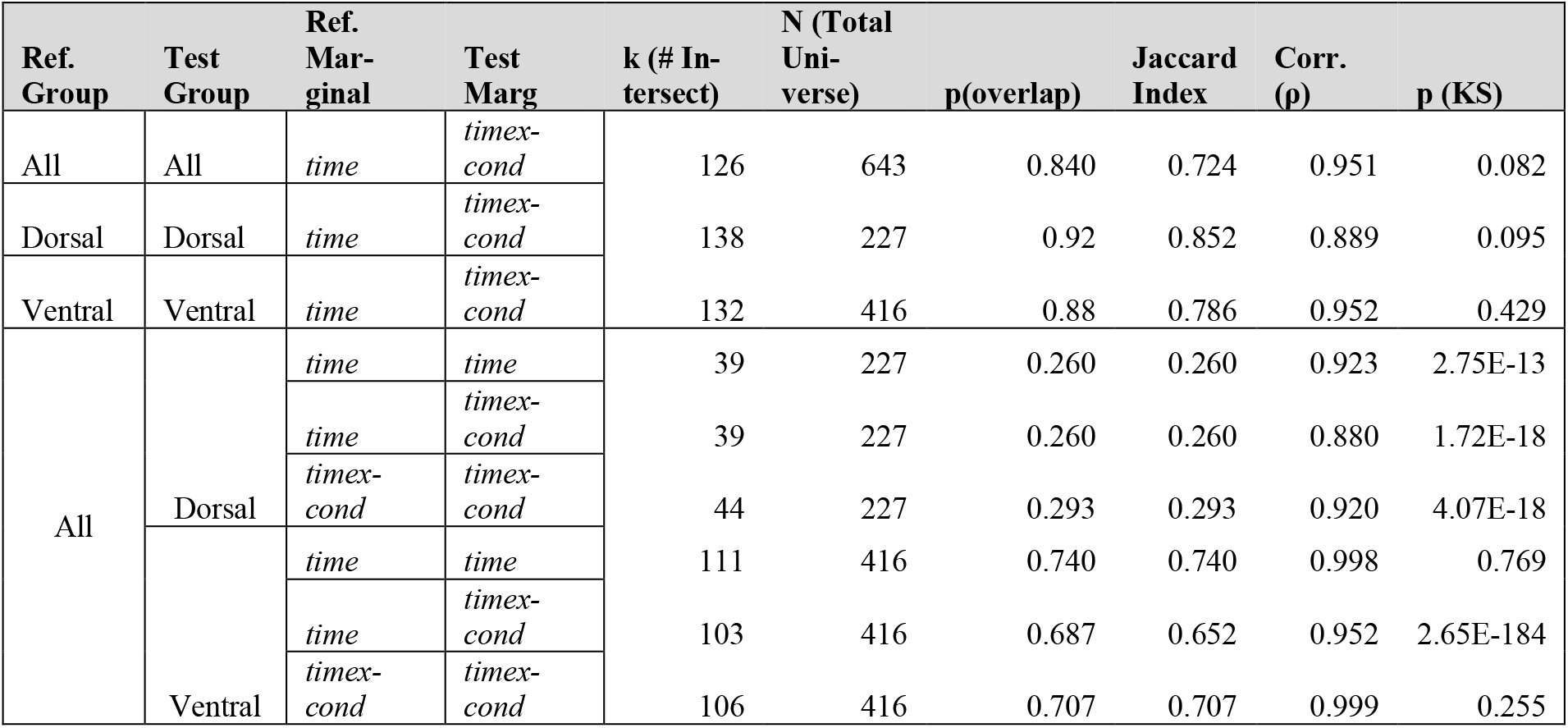
Tests of neuron index similarity within marginalizations and across data sets. Here, the intersection of the top 150 contributing neurons to a marginalization (time, timexcond) for each test set was tested against the top 150 neurons for a reference marginal the ensemble (all) group. Encoder weight magnitudes were summed across all components within a marginal to determine composite effect. K = Number of intersect neurons (universe intersection). N = Size of the universe. P(Overlap)= k/N. Jaccard Index = Set intersection/Set Union. Spearman Rho = Correlation in the encoder weights of the top 150 neurons in each group. p(KS) = 2-Way Kolmogorov-Smirnov test of encoder weight distribution similarity.

Within the 643-neuron ensemble, ventral neurons were more active than dorsal neurons (**Figure 3G**) (Rank-Sum; W= 567294; N=643; p= 5.28e-07). Overall neural firing rate also varied across the 8 animals (**Figure 3H**; Kruskal-Wallis; H(7,635) =73.0612; p=3.55e-13) (**Sup. Table 3**). While overt patterns in neural activity could not be identified across the recorded ensemble for each rat when sorted by depth (**Figure 3I**), neural responses formed a continuum when sorted by spike rate peak timing (**Figure 3J**), with enrichment during the reaching epoch. This pattern was retained when compiling all neurons across the ensemble (**Figure 3K**), and again when separating the pooled ensemble into the dorsal or ventral neural groupings (**Figure 3L**). Cumulatively, these results indicate that single-unit responses were heterogeneous, generally upregulated during motor behavior, and only weakly structured across conditions (See **Sup. Table 4**), motivating additional population-level analyses (For a fuller description, see Supplemental Material **S3**).

**Table 3:** Dorsal and ventral dPCA component activation optimal phase lags. The optimal lags for time series between each dPCA marginals were determined through cross-correlation of ventral and dorsal components, independently for each component and motor condition (positive = observed dorsal leads, negative = observed dorsal lags). 95% Confidence intervals were calculated via resampling trials (n=1000) to produce PSTH surrogates, then projecting these spike rates through the common observed dorsal or ventral dPCA encoder matrices. Lags were considered consistently non-zero when the bootstrap confidence interval excluded zero. Linear correlations between comparable dorsal and ventral were performed after phase alignment.

| dPCA Marginal | # | Optimal Lag ($V_t - D_t$)<br>Milliseconds | | 95% Confidence Intervals<br>[Lower, Upper] | | Post-alignment<br>Correlation ($\rho$) | |
| --- | --- | --- | --- | --- | --- | --- | --- |
|  |  | Ipsi | Contra | Ipsi | Contra | Ipsi | Contra |
| **Time** | 1 | 0 | 0 | [0,0] | [0,0] | 0.9872 | 0.9883 |
|  | 2 | -9 | -4 | [-24,2] | [-31,-2] | 0.9745 | 0.9665 |
|  | 3 | 34 | 39 | [25,62] | [12,62] | 0.7643 | 0.7195 |
| **TimeXCond** | 1 | -39 | -37 | [-100,-40] | [-60,0] | 0.9783 | 0.9774 |
|  | 2 | 0 | 0 | [-75,0] | [-67,42] | 0.8605 | 0.8750 |
|  | 3 | -6 | -8 | [-40,16] | [-37,18] | 0.9105 | 0.9347 |

### Neural Population Analysis

#### Ensemble Data Occupies a Low-Dimensional State Space (PCA)

We next asked whether the entire neural ensemble (e.g., **Figure 3J**) could be used to extract a smooth latent state space during reach. Principal Component Analysis (PCA) of trial-averaged neural firing rates within the binned data (−500 ms to +1000 ms around movement onset) revealed a strong low-dimensional structure across the ensemble: Within the full dataset, the first six PCs explained ∼91% of total variance (PC1–PC3 ≈ 39%, 24%, 17%; PC4–PC6 ≈ 5%, 4%, 2%, respectively). This low-dimensional structure suggests that neural population activity is effectively governed by a few shared processes and justified focusing subsequent dynamical analyses on a small set of latent axes.

#### Neural Components show Shared and Divergent Neural Modes (jPCA)

To examine whether the low-dimensional population activity exhibited dynamical structure signatures, we applied joint principal components analysis^2^ (jPCA) to the trial-averaged spike rates (Methods). Because jPCA first performs PCA and retains the top 6 components, then finds an optimal rotation of this 6-dimensional basis, our jPCA results collectively explain the same 91% ensemble variance as our 6-dimensional PCA. We found three temporal motifs within the three orthogonal jPCA planes: 1) a rotational mode [JP1], 2) a transient ramp-like mode [JP2], 3) a smaller condition-divergent mode [JP3].

##### [JP1] A shared rotational component aligned to the reach epoch (*Fig 4A*)

The first jPC plane exhibited a clear rotational trajectory during the reach (**Figure 4A**), forming an ellipse in state space whose direction and amplitude were similar for ipsilateral and contralateral reaches. This plane accounted for 13% of jPCA-captured variance and resolved mostly within the 0-250 ms interval (**Figure 4B**), which corresponded to the average reaching duration (ipsi: 212 ± 109 ms, contra: 249 ± 61 ms).

**Figure 4:**
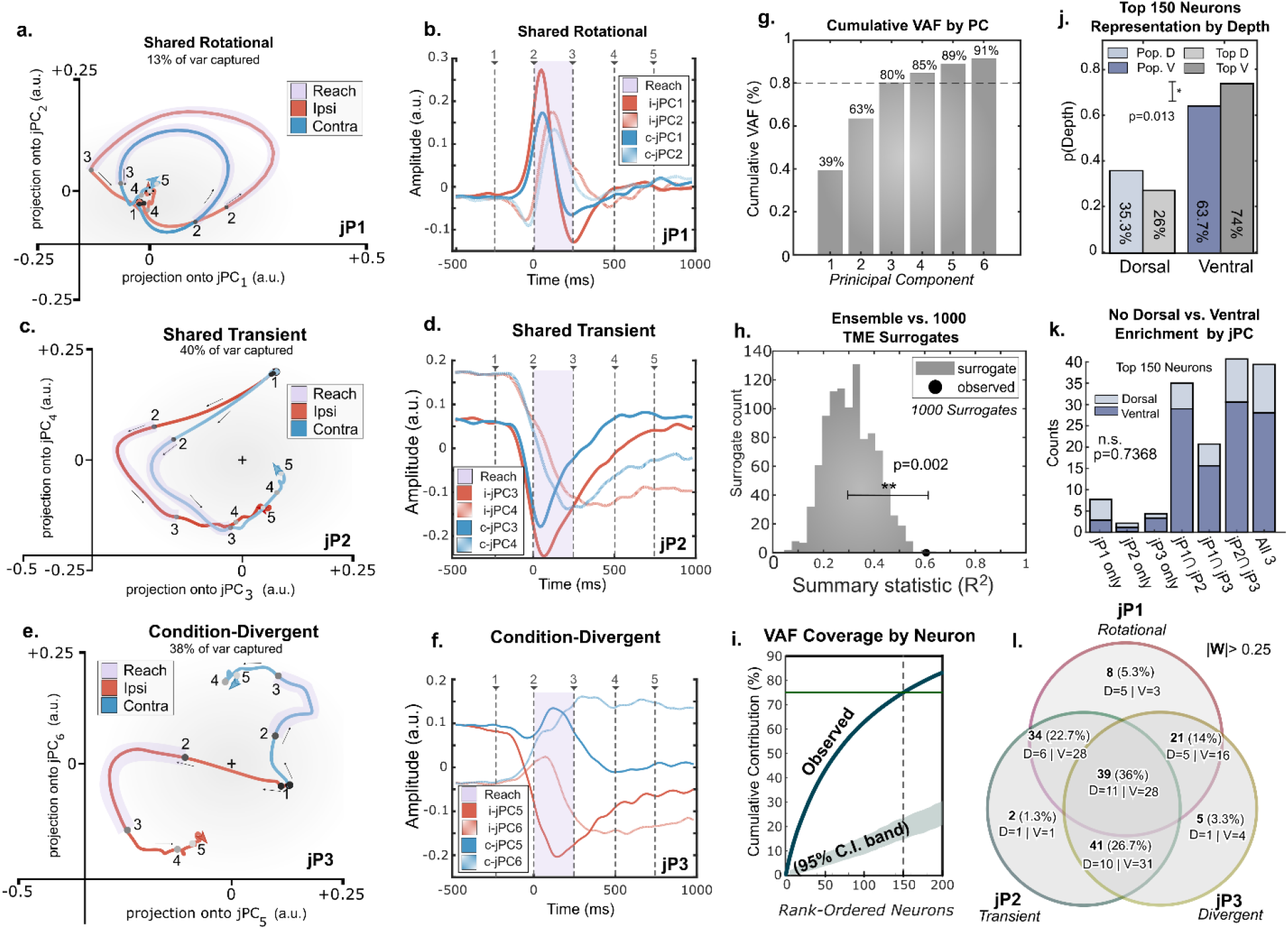
Ensemble data is well-represented by a low-dimensional state space. A) jPCA plane 1 corresponds to an rotational trajectory between two components. Here, time evolution is ordered by numerical sequence. B) Time series representation of jP1 for ipsilateral (i) and contralateral (c) reaches. Numbered time points correspond to traces given in A. The usual reach-active epoch (0-250 ms) is overlaid in purple. C) jPCA plane 2 corresponds to a transient component, numbered as in A. D). Time series representation of JP2. E) Condition-divergent jPCA plane 3 component F) Time series representation of JP3. G) Cumulative variance accounted for by principal component on 643-neuron ensemble across both conditions. H) R2 statistic of the observed skew symmetric matrix of the ensemble data as produced by jPCA vs. 1000 shuffled surrogate tensors generated by tensor maximum entropy (TME) resampling. I) The contribution of individual neurons to PCA cumulative variance accounted for. The 95% Confidence interval band, assuming purely random weighting is overlaid in green. J) Representation of dorsal/ventral neuron percentage in the top 150-weighted neurons vs. the entire ensemble. K) Neurons whose normalized projection-weight magnitude was >0.25 were considered to contribute to plane, and could contribute to multiple planes. Count of top-weighted 150 dorsal and ventral neurons by their contributions to the three jPC planes, or combinations thereof. L) Venn-diagram of the counts of the top responding neurons to the three jPC planes, or combinations thereof.

##### [JP2] A robust shared transient component reflecting movement progression (*Fig 4B*)

The second jPC plane showed a transient trajectory which described the largest share of jPCA-captured variance (40%, **Figure 4C**). This transient component rose during movement initiation but did not return to baseline by the end of the analysis window (**Figure 4D**). This component captured a stereotyped temporal progression through the reach, consistent across conditions and animals.

##### [JP3] A motor condition-divergent component

The third jPC plane captured the divergence between ipsilateral and contralateral reaches, emerging shortly after movement onset (**Figure 4E**). This plane represented 37% of jPCA-captured variance, and was the only jPCA plane with appreciable condition differentiation in neural trajectories. Divergence between the ipsilateral and contralateral neural activity increased during the motor-active period (**Figure 4F**).

Together, these three components describe a dominant temporal progression and limited condition-specific divergence. Ensemble neural data could be captured by such a low-dimensional description (**Figure 4G**) significantly better than could be expected by chance (**Figure 4H**), supporting application of these motifs.

#### Neurons in the dorsal and ventral zones contribute similarly to dynamics motifs

We next evaluated how individual neurons contributed to these motifs by considering the top 150 neurons (corresponding to 78% VAF) by contributing weight in jPCA space (**Figure 4I**). While exact numbers shifted by preprocessing parameters, we found that the majority of the most influential neurons (135/150, 90%) consistently contributed to more than 1 jPC plane. We then tested the cooccurrence of neural contributions of the top 150 contributing neurons to jPC planes 1-3 (defined as *w* > ‖0.25‖ for the unity-normalized weight matrices). We observed 111/150 (74%) of the top neurons were from the ventral partition, representing a significant enrichment relative to the ensemble (two-tailed binomial test, prob(111)=0.647; p= 0.013) (**Figure 4K**). However, there was no evidence of neural modularity to these planes [*χ*^2^(df=6, n=150) =7.67, p=0.7368)] (**Figure 4L**). This lack of specialization is consistent with distributed neural contributions to neural dynamical motifs rather than distinct functional subpopulations.

#### Dorsal and ventral neurons express the same jPCA motifs

We next further considered whether these recovered dynamics motifs differed across depth (**Figure 5**). Thus, we performed jPCA separately on dorsal and ventral populations. When jPCA was applied independently to dorsal (n=227) (**Fig 5A-F**) and ventral (n=416) (**Fig 5G-L)** neuron groups, these analyses recovered the same dynamics motifs as the jPCA on the ensemble.

**Figure 5:**
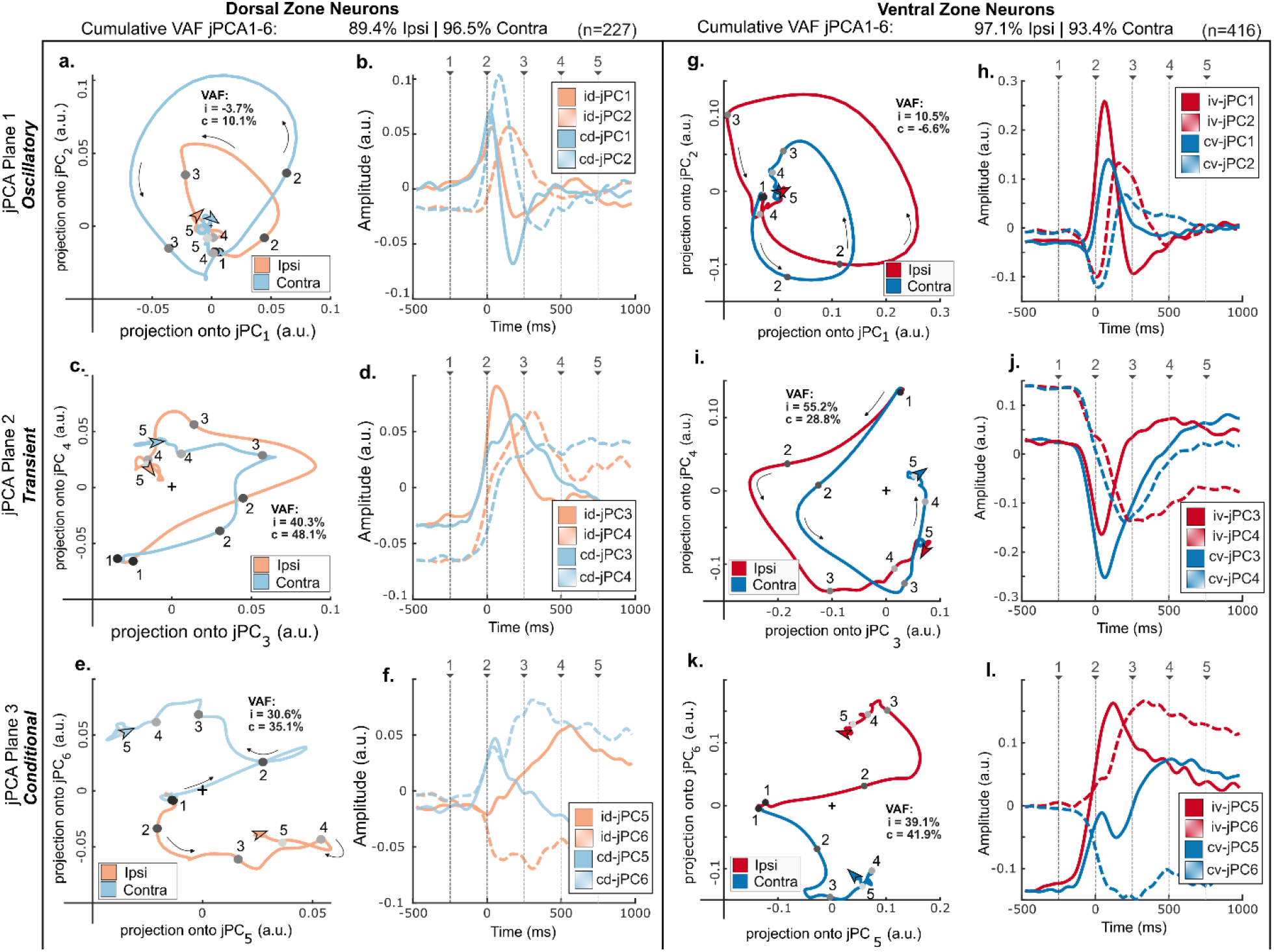
Neurons from the dorsal zone (n=243) and ventral zone (n=435) were analyzed separately using jPCA. jPCA extraction prioritizes planes which maximize rotational dynamics. In both cases, the same three dynamics motifs are extracted as with the neural ensemble: 1) a rotational mode, 2) a transient mode, 3) a reach-conditional mode. The explanatory power of each mode was independently measured by back-projecting spike rates from jPC planes, and calculating explained spike rate variance accounted for (VAF). For both dorsal and ventral components, the transient component (jP2) provided the most explanatory power, with similar timing for the ipsilateral and contralateral reaching variants. (i = ipsilateral; c = contralateral; d = dorsal; v = ventral)

Both dorsal and ventral jPC analyses showed remarkably high canonical correlations with the ensemble (ρ=0.985-1.00) (**Table 1**). While the distribution of shuffled surrogate statistics was tight and robust for both the transient (jP2) and condition-dependent (jP3) dynamics motifs (95% C.I.: 0.915-0.999), the trial-bootstrapped surrogate distribution variance was much broader for the rotational component (jP1) (95% c.i. 0.594-0.993). This result indicates that while similar transient and condition-dependent jPCA planes could be robustly extracted from across different samples of the ensemble data, the exact rotational plane was more sensitive to the neurons used to derive it (i.e., different combinations of spikes would give different rotational planes). However, individual neurons were generally not as sensitive to specific trial selection when comparing within animals (**Sup. Fig 2**), hence this rotational plane may be more animal-specific (**Sup. Table 5**).

While the shuffled distributions showed a high variance in jP1, both dorsal and ventral neurons showed good alignment with the jP1 extracted across the entire (n=643) neuron ensemble (Dorsal ρ=0.938, Ventral ρ=0.999) (**Table 1**). Notably, the jPC planes derived from the ventral neurons were statistically more correlated with the jPC planes from the ensemble than nearly all the depth-shuffled null statistics (i.e., an admixture of ‘dorsal’ and ‘ventral’ coded neurons). Despite the larger ventral sampling population, dorsal and ventral subspaces remained highly aligned (See also **Sup. Table 6**). Our jPC motifs are therefore not depth-specific.

#### Common neural dynamics are dominated by shared time-varying marginal components (dPCA)

Our jPCA analyses indicated a similar dynamical structure across depths and reaching variants, with a rotational component. Population-level similarity, however, does not preclude condition-specific coding at the level of individual neurons. Indeed, we observed that ∼23% of neurons were active in only a single condition, indicating substantial condition-specific variance that a shared population-level description alone may not capture. To isolate the putative shared time-dominant component from the condition-specific component, and to determine whether these two forms of coding are segregated across the neurons or populations, we applied demixed principal component analysis (dPCA)^9^ (**Figure 6**).

**Figure 6:**
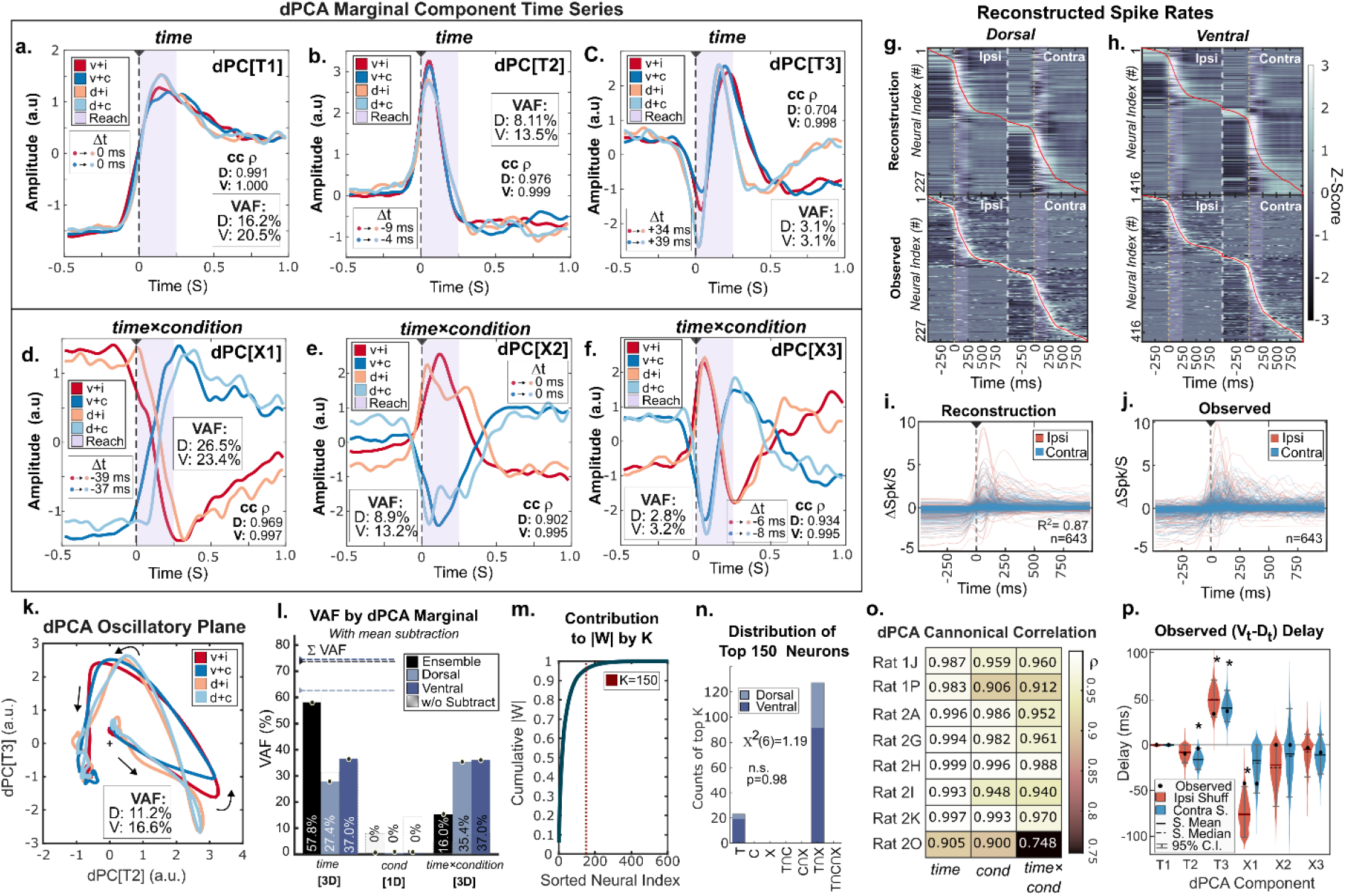
Demixed Principal Component Analysis (dPCA) marginalizations. dPCA consistently extracted three components across the *time* marginalization **A-C**) between independent samples of the ensemble (i.e., dorsal (light) or ventral (dark) neurons). When mean-subtracting neurons, three *time×condition* interaction components were also extracted (**D-F**). **G-H**) Reconstructed, sorted, spike rates from the dPCA components (top) vs. the observed spike rates (below) for each neuron (row) displayed as a heatmap. Red = The peak timing of the neural PSTH. Sort order for the reconstruction was defined from the observed data. **I-J**) Reconstructed vs. observed spike rates for the retained ensemble (n=643). **K**) The pair of T2 and T3 *time* components could describe a rotational dPCA plane, as with jPCA. **L**) The variance-accounted-for (VAF) of each dPCA marginal (black = ensemble, teal = dorsal, blue = ventral). The VAF without mean subtraction is overlaid for each marginal as a dashed box. **M**) The cumulative contribution of most-influential neurons on dPCA (ranked by encoder weight magnitude). **N**) The distribution of the top 150 neurons, independently drawn from the dorsal and ventral neural populations. **O**) The canonical correlation of dPCA marginals of each rat to the ensemble. **P**) Confidence intervals of delays between dorsal and ventral components were identified by bootstrapping over trials. Components whose confidence intervals did not include zero were considered to have significant delays. (v.i. = Ventral Ipsilateral; v.c. = Ventral Contralateral; d.i. = Dorsal Ipsilateral; d.c. = Dorsal Contralateral) The average reaching epoch, 250ms, is overlaid in purple.

Here, dPCA decomposed population activity into three marginalizations across neural activity coded by reaching condition (ipsilateral/contralateral lever): *time*, *condition* and *time×condition* (i.e., common components varying over time, primarily shared between reaches; components with positive/negative sign differing between ipsilateral/contralateral reaches; and time-varying components whose sign depended on reach condition). We decomposed neural activity into the sum of the top 6 total components. Because our mean-subtraction preprocessing step trivialized *condition* components, we extracted 3 components for the *time* marginal (T1-3) and 3 components for the *time×condition* marginals (X1-3).

With our dPCA parameters established, we applied dPCA independently to the ensemble, dorsal, and ventral neural spike rate datasets to determine which marginalization contained the similar dynamical structure observed in jPCA (**Figure 6**). When utilizing the ensemble, we found that the 3 components of the *time* marginalization largely captured activity across the neural ensemble, accounting for 57.8% of the variance. Strikingly, when dPCA was run independently on dorsal and ventral (**Fig. 6A-C**) subpopulations, and prior to any alignment, each group exhibited nearly identical temporal components, (canonical correlation 0.9953-1.0). Interestingly, the three *time×condition* marginals captured more of the variance when dorsal and ventral populations were analyzed independently (Ensemble 16.0%; Dorsal 35.4%, Ventral 37.0%) (**Fig. 6D-F, L**), indicating that motor-conditional differences in neural activity are more prominent within depth regions than across it. To reconstruct this neural activity, we derived expected spike rates via back-projection of dPCA marginal component drives. Reconstructed neural spike rates grossly fit observed spike rates during the motor active period in the dorsal (**Fig 6G**) and ventral (**Fig. 6H**) partitions along with the ensemble (**Fig 6I-J**), indicating dPCA captured an effective representation of neural data (R^2^=0.87).

We also made the incidental observation that dPCA *time* marginals T2 and T3 together form a rotational-like plane, which resembles a similar structure observed in jPCA (**Fig. 6K**). Interestingly, the variance accounted for by the putative dPCA plane (Dorsal: 11.2%, Ventral: 16.6%) is comparable to the rotational element observed in jPCA plane (10-13%). Indeed, *time* marginals were most robustly extracted across rats (**Fig 6O**) (alignment to ensemble all 0.905-0.999 canonical correlation), although Rat 2O *time×condition* marginals fit considerably worse than other rats (c.c. ρ=0.704).

#### dPCA marginals are broadly encoded by overlapping sets of neural components

As with jPCA (**Fig 4A-F**), we next considered if different dPCA marginals (*time, time×condition*), originated from the same neurons, or were differentially organized by depth. We ranked individual neurons by their contributions to these marginals by summing over the magnitude of each neuron’s per-marginal encoder coefficients, normalized to each neuron’s total magnitude (**Fig. 6M**). As most of the variance accounted for could be captured within the first 150 neurons (>95%), we again considered this population as ‘most influential’ for determining neural contributions. Within the top 150 most influential neurons to dPCA marginals, ventral neurons were slightly enriched (73.3%) relative to sample ratio (64.7%) (Binomial Test, prob(110)=0.647; p= 0.0207). However, as with jPCA, we found no difference between the distribution of marginal contributions of the top 150 neurons by dorsal or ventral identity ([*χ*^2^(df=6, n=150) =1.19, p=0.9773)] (**Fig. 6N**).

Because most of the top 150 most influential neurons contributed to both *time* and *time×condition* marginals, we also independently ranked the top neurons for *each* marginal and evaluated the size of the intersection between marginals and across depth groups (**Table 2**) (e.g., ‘how many top neurons for the ventral *time* group were also top neurons in the ensemble *time* marginal?’). Consistent with prior results with jPCA planes, the overlap between dPCA *time* and *time×condition* marginals were substantial *within* the ensemble, dorsal, and ventral dataset (Jaccard=0.72-0.85). A similar overlap was found between top contributing neurons to the same marginals *across* the dorsal/ventral groups and the ensemble (Dorsal↔Ensemble: Jaccard=0.24-0.29; Ventral↔Ensemble: 0.60-0.74). Thus, as with the jPCA planes, neurons simultaneously carry the time and condition information within the dPCA marginals, (**Fig. 6N**), rather than forming segregated dorsal or ventral subnetworks of specific neural units.

#### Common dynamics occur with both synchronous and asynchronous timing

Having observed that the dPCA components were nearly identical in the dorsal and ventral partitions, and broadly distributed across the neurons, we next considered whether these components were expressed with the same timing in the dorsal and ventral populations (**Fig. 6P**). We evaluated the phase relationships by cross-correlating the time series between like-components of the dorsal and ventral-derived dPCA to identify the optimal lag, independently for both reaching conditions (**Table 3**). We estimated confidence intervals from the distribution of equivalent lag measurements across 1000 trial-bootstrapped surrogate datasets, whose spike rates were projected through the data-trained dPCA encoder matrices.

No significant differences of optimal lags were sustained between like components of ipsilateral and contralateral reaching behaviors after Bonferroni correction (Monte Carlo Permutation Test (Two-tailed, p=0.046-1.0). However, significant differences between different components of the ipsilateral (Kruskal-Wallis, H(5)=4808.977, p=0) and contralateral (Kruskal-Wallis, H(5)=3369.62, p=0) reaching behaviors were retained. *Post-hoc* comparisons of different components, within reaching condition, determined that all components significantly differed in their optimal lag alignments (All components p<1.05e-17, Tukey Corrected). Two exceptions were noted for the contralateral reaching condition only: *time* dPC-T2 and *time×condition* dPC-X1 components showed no difference (p>0.999), nor did dPCA-X2 and dPC-X3 (p=0.9686).

Were observed lags meaningful? We considered an optimal lag to be consistently non-zero when the trial-bootstrapped confidence interval excluded 0 ms (α=0.05). Two drives returned as consistently non-zero: 1) the low-variance *time* component dPC-T3 (See supplemental S4.6) and 2) the (ipsilateral) condition-specific dPC-X1 drive. Lags between dPC-X2 and dPC-X3 were not consistently non-zero, but had a larger range of delays across bootstraps. The predominant *time* component, dPC-T1, represented an extreme case, where both the dataset and *all* 1000 surrogate samples returned 0 phase differences between dorsal and ventral components (**Table 3**), strongly supporting synchrony of this element across neural depth and motor conditions.

Our results cumulatively indicate spinal neural dynamics 1) may be shared across dorsal and ventral populations, 2) manifest with little evidence of neural specialization or modularity, 3) may display components whose time courses are simultaneous, phase-lagged, or depth-dependent.

## Discussion

### Summary of findings

The principal findings of this paper are threefold. First, we have pioneered a method for high-density recording of cervical spinal neural activity during forelimb movement in awake, behaving animals. Second, we confirmed the presence of the neural population dynamics, primarily observed in the cerebral cortex, in the cervical spinal cord during voluntary forelimb movement. Third, the dominant population dynamics were conserved across both movement conditions and spinal depth. We discuss each of these points below.

### Novelty of Awake Cervical Spinal Recordings in Rodent

Although comparable approaches for recording neural activity during awake behavior have been established in the lumbar cord^11,12^, the cervical spine serves different biomechanical functions than the lumbar spine^13^. The cervical vertebrae are surrounded by thick paraspinal and nuchal musculature^14^, and are considerably mobile^15^, while the cervical spinal cord also moves more rostro-caudally than lumbar cord^16^, and during cardiorespiratory events, complicating neural signal recording quality^17^. Consequently, stabilization for recording must span multiple vertebrae. Thus, while studies in reduced^4,7^ or anesthetized preparations^6,18^ existed, there had been no successful high-density neural recordings from the cervical spinal cord in awake, behaving animals. Despite these challenges, recordings were stable in our preparation, with electrode drift confined to ±50 μm (**Figure 2A**), allowing us to compare neural dynamics across spinal depth.

The spinal cord separates the inputs and outputs of sensorimotor transformation along the dorsoventral axis: Primary afferents generally terminate dorsally, while motor output is generated ventrally. Resolving this axis was essential to our study, thus we exploited the linear contact arrangement of the Neuropixels silicon probe to assess depth. Channels were assigned using the polarity and latency of the local field potentials evoked by radial nerve stimulation^19^, with the dorsoventral border placed at the sign reversal occurring 1–3 ms after stimulation (**Fig. 2C-D**). Our depth-profiling method thus allowed us to simultaneously sample across dorsal and ventral lamina of the spinal cord with a single probe. This is an advantage compared to cortical areas, which would generally require multiple probes to record from the comparable sensory-associated S1 and motor-associated M1 regions.

This method also allowed us to evaluate probe depth online and reposition until a comparable profile could be attained in each animal. Combined with fixed stereotaxic entry points and insertion depths, our method should reduce inter-animal variability in cervical spinal recording profiles.

### The Observation of Neural Dynamics in the Cervical Spinal Cord

Low-dimensional neural population dynamical motifs have been described in sensory^20^ and motor cortices during reaching^2,3^ and grasping behaviors^21^, and been attributed to both premotor planning and execution phases of motor control^22^. Comparable low-dimensional structure has since been reported in the lumbar cord, where rotational dynamics accompany rhythmic locomotor output^4,7^. Our results extend these observations to the cervical cord during skilled reaching.

Unlike locomotion, reaching is a discrete, non-repeating movement^23^ with no periodic drive to inherit, and varies in both direction and motor pattern across different targets. As afforded by our preparation, here we identified that neural activity in the cervical spinal region, like the cortex, can also be described by low-dimensional neural dynamics. This indicates that such activity patterns represent a general property of neural activity across the neuroaxis, rather than residing only within the cortex or manifesting as a byproduct of rhythmic motor behaviors within the spinal cord.

### Consideration of Cervical Neural Dynamics by task, timing, and depth

#### Comparison of neural dynamics across depth

Using jPCA and dPCA analyses, we extracted neural dynamics patterns across all recorded cervical spinal neurons, and then independently for dorsal and ventral neurons. Generally, we observed that the patterns extracted across the ensemble matched those extracted from the dorsal or ventral patterns independently (**Fig. 4-6**). This similarity was surprising given the dorsoventral anatomical specialization of the cord for receiving afferent input vs. organizing efferent output, and suggests that the recovered dynamics do not uniquely reflect these roles. Neural dynamics are patterns of activity extracted across the population: Similarity in observed neural dynamics across depths and behavioral variants suggests that diverse descending, sensory, and local influences converge within a common low-dimensional population structure.

#### Comparison of neural dynamics by motor behavior

Some neural dynamics patterns were not only similar across depths, but across motor behaviors. In our task paradigms, the animal uses the same limb to commonly reach forward, then towards a left or right target: Commonalities in some aspects of the neural dynamics are thus unsurprising. While the dPCA method seeks this cross-conditional component with the *time* marginalization, jPC1-2 also returned this common structure implicitly. In either analysis, these common components contained considerable explanatory variance (jPC1–2, 53% of jPCA-captured variance; dPCA *time* marginalization, 57.8% of ensemble variance) across behaviors.

Nonetheless, the presence of a substantial component distinguishing between reaching targets (jPC3, 37% of jPCA-captured variance; dPC-X1–3, 16% of ensemble variance and 35–37% when dorsal and ventral populations were analyzed separately) indicates that our dynamics are not limited to generic movement-related signals, but also carry information about the particular movement being performed. These condition-specific dynamics could reflect spinal segmental differences in the sensorimotor preparation and execution of different reaching behaviors.

#### Relative timing of neural dynamics by depth and condition

Our dPCA results produced patterns which were clearly similar across depth. To test whether this similarity also reflected synchronous dorsal and ventral dynamics, we performed a lag analysis. We noted two observations in the data: The dominant *time* component (dPC-T1) was synchronous across depth, whereas the dominant condition-dependent component (dPC-X1) showed dorsal-ventral delays.

First, the dominant *time* component (dPC-T1) was precisely synchronous across depth: The dorsal-ventral lag was zero (at 1-ms resolution) in both reaching conditions, and in all 1000 trial-bootstrap resamples. Such synchrony is difficult to reconcile with serial propagation from one region to the other, and instead suggests a simultaneous common drive within dorsal and ventral populations. Because dPC-T1 was also shared across reaching conditions, such a drive could feasibly be observed in the spinal cord during other phasic movements of the forelimb.

Second, the dominant *time×condition* component (dPC-X1) showed a contrasting pattern. In the ensemble and depth-segregated data, the ventral component *preceded* dorsal component by approximately 40 ms in the point estimates of both reaching conditions (−39 and −37 ms; Table 3). Estimations of this ventral-to-dorsal lag were more unstable than dPC-T1, as only reaches towards the ipsilateral target had a bootstrap lag confidence interval which excluded 0 ms. However, if sustained, this lag exceeds the latency of the entire segmental reflex arc in the rat forelimb (∼5–6 ms,^24^), and is instead more comparable to the interval by which rat forelimb muscle activation precedes movement onset during reaching (∼30–60 ms^25^). This timing is compatible with a reafferent contribution to the cervical spinal dynamics, in which condition-specific ventral motor output generates condition-specific sensory feedback subsequently in the dorsal population.

Together, the coexistence of a synchronous condition-invariant component and a ventral-leading condition-dependent component suggests that cervical spinal dynamics integrate at least two influences, such as a common drive and movement-related feedback, rather than arising from a single homogeneous process.

#### Relationships between single neurons and neural population dynamics

Neural population dynamics are derived from the coordinated activity of individual neurons. Single unit analyses have previously been useful for identification of circuit-level specificity, neural fine-tuning, and modular activations to the motor drive (e.g.,^18,26–29^). Here, we observed that organization emerged as the scale of analysis increased: Individual neurons displayed depth-dependent differences, heterogeneous firing rates, timing, and condition-dependent activity (**Figure 3**), yet population activity nonetheless converged onto a low-dimensional structure which was highly similar across depth, motor conditions, and animals (**Figs. 4–6**).

The single-unit and population-level observations are not contradictory. Population dynamics describe the activity patterns shared across neurons (including both condition-invariant and condition-specific components), whereas single-unit analyses capture how individual neurons deviate from these shared patterns. As neural dynamics analyses prioritize common patterns in activity across neurons, unit-level features and specializations may be lost—relegated to lower-variance latent factors and neglected in consideration. Similar dissociations between neuron-level heterogeneity and similarities in population-level dynamics have previously been reported in both biological^30^ and artificial^31^ neural systems. This disparity reflects the differing scales of analysis: Analyses focused on single neurons or cell types would miss the shared population structure, whereas population dynamics alone cannot explain cell- and circuit-specific processing. Understanding spinal motor control will therefore require both perspectives.

### Potential biological interpretations for observed neural dynamics patterns

The discovery of population dynamics in the cervical cord motivates consideration of its origin and function. In the cortex, neural dynamics have been variously attributed to structured inputs, feedback^32^, and a common timing across behaviors^33^, thereby coordinating descending drive. If low-dimensional population dynamics in the cortex simplify^34^ aspects of top-down control for voluntary movement, one possibility is that spinal dynamics are simply inherited from cortical dynamics through descending pathways. However, the spinal cord is not a passive receptor of descending inputs: The spinal interneuron network translates descending commands and conveys them to motoneurons, while sensory feedback arising from movement is likewise processed and utilized in the spinal cord before being carried to the cortex. These bidirectional interactions between cortical and spinal circuits raise the possibility of bidirectional influences within the neural dynamics as well, and the spinal structure we describe here may also be neither purely inherited from nor independent of its cortical counterpart.

Our depth-resolved recordings allow the role of neural dynamics to be approached from a different angle, because the spinal cord segregates its inputs to outputs along the dorsoventral axis. In the rat, corticospinal commands primarily targets the dorsal horn^35^, as does afferent feedback, while motor output is generated ventrally. The interneuron network linking these two regions can also generate structured population dynamics^4,7^. If spinal dynamics were inherited from a distinct source or reflected stage-specific processing, they should differ between dorsal and ventral populations. However, despite clear dorsal-ventral differences in single-neuron activity, population dynamics were highly similar across depths. Spinal neural dynamics during reaching thus appear to combine descending drive with locally generated and feedback-driven activity in a manner that produces similar activity across the cord. This suggests that the observed dynamics do not primarily reflect input- or output-stage-specific processing in the spinal cord, but instead reflect a structure shared across these regions. We consider two non-exclusive interpretations of this finding below.

One interpretation is that the observed spinal population dynamics encode temporal structure that is partially independent of specific sensory or motor representations. Similar ideas have been proposed for locomotor (e.g.,^36^) or other spinal motor behaviors (e.g.,^37^), where spinal activity may provide coarse timing information for diverse spinal circuits. Accordingly, the condition-invariant component of the spinal population dynamics observed here may reflect a common timing ‘scaffold’ for behavior that need not contain the specific transformation required to reconstruct muscle activations. In this view, differences in motor pattern between reach targets would arise not from changes in the scaffold itself, but from which circuits are engaged along it; for example, which motor pools and premotor interneurons are recruited, or which segmental reflex pathways are gated, at each phase of the shared time course.

A second, complementary, interpretation is that dorsal and ventral dynamics are similar because sensory and motor processing are strongly coupled in the spinal cord: Reaching to different targets not only varies motor neuron activations, but also sensory feedback thereof. Neural dynamics may reflect circuit structure and constraints^38^: If dorsal and ventral lamina are sufficiently interconnected, network structure could coordinate common activity across the cord (as previously observed *in silico*^39^). Candidate neurons mediating this communication across spinal depths include those mediating the spinal reflexes, sensory feedback^40^, or autogenic drive^41^. By extension, control of different motor behaviors by descending drive may shape these shared dorsal-ventral neural dynamics by modifying motor activity *or* filtering sensory drive, providing additional, non-intuitive mechanisms for control.

## Conclusions

We developed a new paradigm for recording from the cervical spinal cord of the awake rat, enabling, for the first time, simultaneous depth-resolved characterization of spinal population dynamics during voluntary reaching behavior at this scale and resolution.

Spinal population activity was low-dimensional and shared across depths and behavioral conditions. Neural contributions to these dynamics were distributed across depth groups, and showed no evidence of modular organization. Within this shared structure, the dominant condition-invariant component was highly synchronous across depth, whereas the dominant condition-dependent component showed a variable, ventral-leading, delay consistent with timescales expected for movement-related feedback.

Together, these findings position the spinal cord within a broader framework of computation through population dynamics, extending principles previously identified in cortical circuits to a downstream motor structure. More broadly, these results suggest that anatomical specialization and population organization may operate at different scales of description: While dorsal and ventral neurons differ substantially at the level of individual responses, their ensemble activity evolves along a common low-dimensional temporal scaffold.

Finally, the experimental and analytical framework developed here, including depth-resolved Neuropixels recordings, provides a foundation for probing spinal population dynamics in the cervical spinal cord with high spatiotemporal resolution. Future work combining perturbation, multi-site recording, and cell-type–specific analyses will be necessary to resolve the causal origins of these dynamics, their relationship with cortical dynamics, and their functional role in motor control.

## Methods

### Animal and Task Model

Eight adult male Long–Evans rats (350–430 g, 10–12 weeks old) were trained to perform a skilled reach-to-lever task for water reward. To allow us to compare neural responses across distinct motor conditions, we trained animals to reach with the right forelimb toward one of two lever positions, placed ipsilateral or contralateral to the reaching limb. Lever angle, video (240 FPS), and electrophysiology were synchronized via TTL events. The hand position was extracted from video recordings (GoPro^TM^ Hero 10 Black, 240 FPS) via DeepLabCut^42^. All experiments were approved by the Experimental Animal Committee of the National Institute of Neuroscience (approval number: 2023028). A fuller description may be found in the supplemental material (**S1.1**, **S1.3**).

### Surgical Methods

Animals underwent two surgeries (**Figure 1A**). First, a custom 3D-printed head plate was attached to the skull for head fixation during behavior. After recovery and reaching training, a second procedure implanted a custom spinal recording chamber (**Figure 1B-D**) spanning C2–T2 vertebrae (see Extended Methods). The animal was allowed to recover for two weeks.

During acute recording, a ground screw was inserted at C7 vertebrae, and a laminotomy at C5 vertebrae allowed probe entry into the right spinal cord. A Neuropixels 1.0 NHP-short probe was lowered into the cervical spinal cord using a micromanipulator interfaced with a removable adaptor. Probe placement was verified online using sensory-evoked potentials (SEPs) generated by percutaneous radial nerve stimulation (0.1 ms pulses), which provided a depth signature allowing coarse assignment of channels to dorsal vs. ventral laminae (see below). Once an ideal depth was reached, the adaptor was permanently fixed to the spinal chamber with UV photocurable resin, and the micromanipulator was removed. A fuller description can be found in the supplemental (**S1.2, S1.4**).

### Depth Labeling vs. Sensory-Evoked LFPs

We sought how neural behavior may vary by motor condition or depth. To evaluate depth, the position of each recording site was estimated using LFP polarity and latency following radial nerve stimulation. Channels exhibiting an early *negative* deflection (1–3 ms) were classified as dorsal^43^, whereas *positive* polarity indicated ventral laminae^19^. Depth labels were verified *post-hoc* using DiI-stained histology of the probe track. When drift occurred during recordings, original depth labels were retained to maintain consistency.

### Neural Preprocessing

Extracellular voltage was acquired at 30 kHz via a Neuropixels base station using Open Ephys^44^ acquisition software. Raw data subsequently underwent high-density motion correction with DREDge^10^ and were spike-sorted using Kilosort 4^45^. Units were manually curated with Phy2. Clusters were further excluded if they exceeded 5% false discoveries (ISI-based estimate) and >0.5% refractory violations^46^. (See **Supplemental Fig 1**, **S2.1**).

A total of 797 neurons were identified across animals, and 643 trial-active neurons were retained for analysis (**Supplemental Table 1**). Spike trains were aligned relative to hand takeoff from home position (time 0). Trials were segmented, and spike trains converted to continuous firing-rate estimates at 1 kHz, and smoothed with a σ=20 ms Gaussian kernel. Neural activity was then binned from -500 ms to 1000 ms relative to movement onset. For neural population activity (PCA/jPCA/dPCA), data were averaged across trials within each condition (ipsilateral, contralateral).

### Tensor Maximum Entropy (TME)

To determine whether observed dynamical structure in the neural data exceeded that expected from the basic statistical properties of the data, we robustly generated sets of 1000 shuffled surrogate datasets using Tensor Maximum Entropy (TME^47^). TME generates data tensor surrogates of a sample which maximize the Shannon entropy of the system, while matching the same features (i.e., first or second-order moments) as the original dataset (iterating over time, neuron, and condition).

We utilized the function *summarizeLDS.m* to calculate the R2 statistic for each surrogate tensor, representing a measure of how much variance could be described via linear dynamical system (LDS). We tested the R2 test statistic from the observed data against the statistics generated by the null distribution of statistics from the TME-surrogates to determine significance^48^ via a 1-tailed t-test, assuming α=0.05.

### Joint Principal Components Analysis (jPCA)

We used publicly available joint principal component analysis (jPCA) code^2^ to evaluate potential rotational dynamics across the neural ensemble. jPCA identifies the skew-symmetric transformation that best captures rotational activity patterns over time. Because rotational structure can arise from sequential activation in neural data^49^, we interpreted jPCA results in conjunction with additional analyses, as described above.

Each neural spike train was baseline subtracted (offset by the average of the first 50 ms) prior to jPCA. Because neurons had variable spike rates and spike density, we did not want the effect of single, rare, spikes to dominate extracted dynamics. Thus, we implemented jPCA with the default soft-norm parameter of k=10, such that spike rates were transformed as 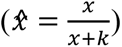. The jPCA default mean-subtraction step removes the cross-conditional mean from the intermediate PCA, calculated from across all conditions. Because we only had two conditions (ipsilateral vs. contralateral reach), this difference is meaningful, therefore we performed jPCA with the PCA mean-subtraction parameter set to false. (Varying preprocessing parameters varied the prominence of the three different jPCA components, symmetry across conditions, or depth, but did not substantially change the primary result. For example, performing the jPCA mean-subtraction step effectively removed the shared transient component [jPC2], without increasing VAF.)

### Demixed Principal Components Analysis (dPCA)

The dPCA technique attempts to separate (‘demix’) the time-varying (*time*), conditional (*cond*), and interacting (*time×condition*) latent factors within the data into separate marginal component subspaces^9^. Consecutive components within each marginal are orthogonal, successively extracted within each subspace (marginal), along the axis of maximal variability (as with traditional PCA). Thus, dPCA method is well-suited for extracting time-varying latent components which are common across different sampling groups (i.e., ipsilateral vs. contralateral reach), and isolating elements which are condition-specific, rather than assigning latent principal components *across* this variability.

Neurons were passed through publicly available dPCA MATLAB code (https://github.com/machenslab/dPCA), regularized using a lambda regularization ladder [λ=0.001, 0.01, 0.1], stepped as necessary to avoid runtime errors. We applied a per-channel, per-condition, mean-subtraction for neural spike rates, storing this offset for signal reconstruction. Due to this mean subtraction step, the dPCA *condition* marginal was trivial, captured by a static offset. (For fitting performed without this mean subtraction, see supplemental **S4.5, Sup. Table 7**.) We used the following number components per marginalization across the ensemble as *time* = 3, *condition* = 0, *time×condition* =3. We then consistently used this same number of components when performing dPCA across different subsets of the same ensemble (i.e., per-rat, per-depth).

To measure the significance for lag testing, we bootstrapped trials to produce 1000 surrogate datasets. Because each trial contained only a single motor condition, during trial bootstrapping, shuffling indices were generated independently within ipsilateral/contralateral reaching conditions to ensure condition-balanced sample depth. Surrogate datasets were projected through the observation-trained dorsal/ventral dPCA encoder matrices to maintain consistent dynamics, then we measured the lags between like components of the dorsal and ventral partitions, independently for the ipsilateral and contralateral reaching conditions. When projecting shuffled PSTHs using the dPCA encoder matrices trained on the ensemble or dorsal/ventral partitioned datasets, shuffled PSTHs were pre-processed as with the original dataset. To avoid spurious correlations outside physiological relevance, we clamped the maximal delay value for cross-correlation to ± 100 ms.

### Subspace Alignment

We compared subspaces (planes: jPCA; marginal components: dPCA) using Procrustes alignment and canonical correlation, quantifying similarity to the full ensemble. When fitting subspaces, the full ensemble was used as the reference against which other groupings (e.g., per-rat, per-depth) were aligned. Because the number of neurons was not consistent by group ID, we generated surrogate data for testing significance of each group separately.

For each alignment test, we had five sets of data:

1. <u>The Observed Ensemble</u>: The full neural ensemble, treated as ground truth.
2. <u>The Groupwise Dataset</u>: Subsets of the full ensemble, grouped according to statistical test (e.g., by rat, by depth).
3. <u>The Group-Complement Dataset</u>: For each element of the group used in #2, we considered the effect of *removing* that group from the dataset (i.e., ‘jack-knifing’).
4. <u>Groupwise Surrogate</u>: Using the same numbers of neurons as the groupwise dataset, we drew 5000 surrogates from the ensemble data. (*This was the null data used to test the significance of grouping effects vs. chance*).
5. <u>Groupwise Surrogate-Complement</u>: Using the same numbers as the Group-Complement Dataset, we drew 5000 surrogates from the ensemble data. (*This was the null data used to test the significance of group-removal effects vs. chance*).

We measured fitting quality via the Procrustes fitting residuals, canonical correlation, and axis correlation, relative to the ensemble. Because returned metrics were effectively scalar distance metrics, we assigned significance via 2-tailed comparison of the groupwise observed test vs. surrogate shuffle statistic distributions. P-values for the observed test statistic, *x,* relative to shuffle distribution *X0* were assigned as *P* = ‖*p*(*x* > *X_o_*) − *p*(*x* < *X_o_*)‖. When 1-tailed tests were used, we assigned P-values as *P* = 1 − *p*(*x* > *X*_0_).

The authors used Copilot Chat in early code generation, some aspects of statistical design, and editing the manuscript for readability. All outputs have been reviewed and revised independently. Authors assume all responsibility for the submitted manuscript.

## Accessibility

The code and datasets used in this paper are available upon request, and will be provided online, following review.

## Acknowledgements

We thank Simon Giszter for the helpful discussions and Moeko Kudo for preparing the histology.

## Supplemental

### S1: Extended Animal Husbandry, Surgical, and Experimental Methods

To enable stable, depth-resolved electrophysiological recordings from the cervical spinal cord in awake, behaving rats, we developed a surgical and experimental preparation combining head fixation, a custom spinal chamber, and Neuropixels-based recording adaptor. This section details the procedures used to achieve stable recordings across dorsal and ventral spinal regions during voluntary movement.

#### S1.1 Animal Husbandry

Eight wild-type male Long–Evans rats (Japan SLC, Shizuoka, Japan; 350–430 g) were used in this study. The experimental procedures were performed at the following ages: habituation (10–12 weeks old), head plate implantation (10–12 weeks old), behavioral training (11–13 weeks old), spinal chamber implantation (14–15 weeks old), retraining (16–17 weeks old), and acute electrophysiological recording (16–17 weeks old). Animals were housed under a standard 12 h:12 h light/dark cycle (lights on from 08:00 to 20:00) in a temperature-controlled environment. Food and water were available *ad libitum*, except before and during the training period (see below). All surgical procedures were performed in accordance with institutional guidelines for animal experiments and the National Institutes of Health Guide for the Care and Use of Laboratory Animals. All experiments were approved by the Experimental Animal Committee of the National Institute of Neuroscience (approval number: 2023028).

#### S1.2 Surgical Methods

We developed a surgical protocol to ensure stable cervical recordings and secure the animal during training. Animals were first habituated to handling (10 min/day for >2 days), followed by head plate implantation. For head plate implantation, animals were anesthetized by intraperitoneal injection of medetomidine-midazolam-butorphanol (MMB^50^) anesthesia (0.17, 1.7, and 2.1 mg/kg, respectively). Once adequate anesthesia was confirmed by the absence of a flexion reflex to paw pinch and by stable pupil size, the animals were positioned in a stereotaxic frame (SR-10R-HT; Narishige, Tokyo, Japan). Anesthesia was then maintained with 0.6–2.5% isoflurane delivered via inhalation for the duration of the surgical procedures. Throughout surgery, heart rate, oxygen saturation, and rectal temperature were continuously monitored and maintained within physiological ranges (heart rate: 180–350 beats per minute; oxygen saturation: >80%; rectal temperature: 35–38 °C). The skin over the skull was incised, and the musculature overlying the bregma and lambda sutures was bluntly dissected. Stainless steel screws were placed in the skull as anchors, and the skull surface was reinforced with a ∼2-mm-thick layer of Super-Bond (SUN MEDICAL, Shiga, Japan). A custom-made 3D-printed head plate (25 × 50 × 2 mm, ∼2.6 g; BioMed Amber Resin; Formlabs, Somerville, MA, USA) was anchored to the skull screws using a UV-curing resin (RelyX Unicem 2 Automix; 3M, St. Paul, MN, USA). Finally, skin around the head plate was sutured shut. The MMB anesthesia was then reversed using atipamezole (i.m. injection).

After a 3-day recovery period, animals were water-restricted for 48 h and underwent lever training (described below). Following an additional 2-week recovery period, animals underwent a second surgery for spinal chamber implantation. As in the head plate implantation surgery, anesthesia was induced by intraperitoneal injection of a medetomidine-midazolam-butorphanol mixture (0.17, 1.7, and 2.1 mg/kg, respectively) and maintained with 0.6–2.5% isoflurane delivered via inhalation. Animals were secured in a stereotaxic frame (SR-10R-HT; Narishige, Tokyo, Japan), and vitals were monitored as described previously. After incising the dorsal skin, the superficial (trapezius and rhomboid), intermediate (splenius cervicis and superficial semispinalis cervicis), and deep (deep semispinalis cervicis and multifidus cervicis) paraspinal muscle layers overlying the C2–T2 vertebrae were sequentially dissected and retracted to expose the vertebrae. The C2 and T2 spinous processes were cleaned with a micro curette to expose the bone and then reinforced with Super-Bond. Super-Bond was also applied to the lateral masses of the C4–C7 vertebrae to facilitate the subsequent fixation of the spinal chamber to the vertebrae during recording.

We developed a custom multipart 3D-printed spinal chamber (**Figures 1A-F**, BioMed Amber Resin, ∼1.3-2.6 g, sized to animal). With the cervical vertebrae fully extended, the chamber base was anchored to the C2 and T2 spinous processes using a UV-curing resin. The larger chamber wall component was then attached to the chamber base and sealed with the same resin. Finally, the chamber clamp post was fixed vertically to enable stable attachment during recording. This configuration balanced the stability of neural recordings with potential compression of the cervical spinal cord within the chamber. The interior of the chamber was rinsed with gentamicin-containing saline and sealed with a silicone adhesive (Kwik-Cast, World Precision Instruments, Sarasota, FL, USA). Finally, the cutaneous tissue was tightly sutured around the chamber.

Animals were closely monitored for 3 days after surgery and received daily intramuscular doses of an antibiotic (ampicillin, 30 mg/kg), an analgesic (ketoprofen, 5 mg/kg), and a corticosteroid (dexamethasone, 0.165 mg/kg). Subcutaneous fluid supplementation and a nutritional gel (DietGel Recovery, ClearH2O, Westbrook, ME, USA; ∼14 g/day) were provided as needed. Animals were allowed to recover for at least ten days after chamber implant surgery before recording.

#### S1.3 Behavioral Methods

After recovering from head plate implant surgery, animals were water-restricted for 48 h and subsequently trained using an operant conditioning paradigm modified from previous studies^51,52^. During behavioral training, animals were briefly anesthetized with sevoflurane inhalation, fixed to a behavioral task system (Task Forcer R1; O’Hara & Co., Tokyo, Japan), and allowed to recover to an awake state before task initiation. Animals were trained to reach with their right forelimb to pull a waterspout lever for a water reward. Each successful reach-grasp-pull movement triggered the delivery of a small water reward (∼7 μL initially) from the spout. To elicit different reaching variants, the lever was positioned either ipsilateral (right) or contralateral (left) to the reaching limb. Animals typically performed an overhand reach for the ipsilateral lever and an underhand reach for the contralateral lever (**Fig. 1H**). During early training (days 1–3), the lever was presented on the ipsilateral side. From days 3–5 onward, sessions were introduced in which the lever was presented on the contralateral side, and an obstacle was placed in front of the left forelimb to restrict its use and enforce task performance with the right forelimb. In later stages (from days 4– 8 onward), animals were trained to voluntarily place their forelimb in a home position 5 cm from the lever before movement initiation. Successful completion of the home positioning–reach– grasp–pull sequence was rewarded with 35–70 μL of water.

Training sessions lasted 30–90 min/day, during which water was provided exclusively (<7 mL/day). When task engagement declined, the obstacle in front of the left forelimb was temporarily removed, and animals were given food to allow a short break before resuming the training session. Animals acquired the task over an average of 8.8 days (range: 4–13 days). Body weight was monitored throughout training to ensure it did not fall below 80% of the pre-restriction baseline. Although supplemental water was prepared if needed, no such intervention was required. After completion of training, animals were returned to *ad libitum* water access and allowed to recover for 2 weeks before subsequent experiments.

#### S1.4 Recording Methods

Prior to the acute recording experiments, animals were reintroduced to the task under water-restricted conditions (2 days) and underwent additional behavioral training for 1–2 days under the same conditions as the final stage of training described above. Each successful trial was rewarded with 35–70 μL of water.

We performed the recording session in three stages:

1. Probe insertion and verification
2. Animal recovery and behavioral recording
3. Post-Recording validation

#### Probe insertion and verification

On the day of recording, anesthesia was induced by sevoflurane and maintained by 0.6– 2.5% isoflurane, delivered via inhalation. Animals were secured in a behavioral task system (Task Forcer R1; O’Hara & Co., Tokyo, Japan), while vitals were continuously monitored as described previously. The silicone adhesive inside the chamber was removed, and a stainless steel screw soldered with stainless steel wire (AS634; Cooner Wire, Chatsworth, CA, USA) was inserted into the C7 vertebra as a ground electrode. Using UV-curing resin (RelyX Unicem 2 Automix; 3M, St. Paul, MN, USA), the ground electrode, previously-applied resin supports on the lateral masses of the C4–C7 vertebrae, and the chamber base were bonded together to ensure mechanical stability. The chamber was leveled relative to the vertebral column by clamping the chamber pole, and a laminectomy was performed at C5 vertebrae using a 0.8-mm burr. To verify the insertion site histologically, a Neuropixels probe (Neuropixels 1.0 NHP Short (NP1015; imec, Leuven, Belgium)) was coated with fluorescent dye (CellTracker CM-DiI Dye (C7000; Thermo Fisher Scientific, Waltham, MA, USA)) for >10 s and air-dried under light-shielded conditions for >5 min. After dural incision with a 29G needle, a pilot hole was created 0.8 mm lateral to the midline under stereotaxic guidance using a 27G spinal needle stylet (UNIEVER 52517600; Unisis, Tokyo, Japan). The Neuropixels probe was then inserted into the spinal cord through this pilot hole at a speed of <2 μm/s using a hydraulic micromanipulator (SM-25C; Narishige, Tokyo, Japan) to a depth of approximately 2.2 mm.

To confirm probe placement, stimulus-evoked local field potentials (LFPs) were recorded by electrically stimulating the radial nerve. A bipolar needle electrode (26G, 3 mm inter-electrode distance) was inserted percutaneously near the lateral epicondyle of the right forearm, and stimulation (pulse width: 0.1 ms; intensity: 10× movement threshold; inter-pulse interval: 0.5 s) was applied. The stimulation site was determined by observing muscle twitches of the wrist and finger extensors following electrical stimulation. LFP signals were amplified (×500) and recorded at 30 kHz using a Neuropixels base station (PXIe-1000; imec, Leuven, Belgium) and Open Ephys software^44^ (v0.6.6; Open Ephys Production Site, Lisbon, Portugal). The ground was connected to the C7 vertebral screw, and the reference electrode was set to the probe tip. Transistor–transistor logic (TTL) pulses were simultaneously acquired via a data acquisition interface (PXIe-1071 and PXIe-6341; National Instruments, Austin, TX, USA) for synchronization of LFP signals with stimulus timing. The depth profile of stimulus-evoked LFPs was evaluated online (Figs. 3C and 3D), and the probe was repositioned if necessary.

#### Animal Recovery and Behavioral Recording

Once an appropriate probe placement was confirmed, the needle electrode was removed, and the probe was secured to the chamber using UV-curing resin, detached from the micromanipulator. The exposed shank was covered with silicone oil (DOWSIL SRX310 Fluid; Dow, Midland, MI, USA). The chamber pole was reclamped to stabilize the cervical posture, and anesthesia was discontinued. For subsequent video analysis, dot markers were applied using permanent ink to the dorsal surface and styloid process of the right wrist joint. Animals were then allowed to recover until fully awake (20–60 min).

For behavioral testing, animals performed the reach–grasp–pull task as described above using the right forelimb. The spout lever was presented either ipsilaterally or contralaterally, and its position was alternated manually every 11 successful trials to minimize potential time-dependent effects. An angular displacement sensor attached to the lever transmitted lever angle to a data acquisition interface (PXIe-1071 and PXIe-6341) as an analog signal ranging from −5 to +5 V. Successful pulling events were identified by threshold crossings at ±3 V. These threshold-crossing events triggered delivery of 35–70 μL of water reward and were treated as the endpoint of successful reaching trials. Corresponding timestamps were recorded as TTL signals via the same data acquisition interface, and a synchronized LED signal was generated simultaneously. Electrophysiological signals were recorded at 30 kHz using a Neuropixels base station (PXIe-1000) and Open Ephys software (v0.6.6) and stored continuously as a single file throughout the session. Behavioral kinematics were recorded using a high-speed camera (GoPro HERO10; GoPro, San Mateo, CA, USA) at 240 FPS with a resolution of 1920 × 1080 pixels, using a narrow field of view (27 mm equivalent) from the right lateral view, with ink markers placed on the wrist joint. Video and electrophysiological data streams were synchronized using the visual (LED) and electrical (TTL) signals.

#### Post-Recording Validation

After recording, animals were re-anesthetized with sevoflurane, and stimulus-evoked LFPs were recorded again using the same procedure to confirm the final probe position, accounting for potential electrode drift during behavioral testing. The Neuropixels probe was then removed for reuse. Animals were deeply anesthetized by intraperitoneal injection of pentobarbital sodium (40 mg/kg) and transcardially perfused with 100 mL of phosphate-buffered saline (PBS; pH 7.4), followed by 300 mL of 4% paraformaldehyde. The cervical spinal cord was extracted and post-fixed for 2 days. The probe penetration site in the spinal cord was confirmed by histological analysis.

Together, these procedures enabled stable, depth-resolved neural recordings across dorsal and ventral spinal regions during voluntary movement.

### S2: Data Preprocessing and Quality Pipeline

To ensure that downstream population analyses were not biased by preprocessing or spike sorting artifacts, we implemented a multi-stage curation and validation pipeline.

#### S2.1 Data Preprocessing

Behavioral kinematics were extracted from high-speed video recordings (GoPro^TM^ Hero 10 Black, 240 FPS) using DeepLabCut^42^ (V2.3.5). An ink mark applied to the dorsal surface of the right wrist joint was tracked during each successful reaching trial. The horizontal position of the mark was numerically differentiated between frames to obtain movement velocity and subsequently low-pass filtered at 30 Hz using a fourth-order Butterworth filter. Initial movement onset was defined as the time point at which the horizontal movement velocity first exceeded 5% of its peak value during the reaching movement and used as time 0 for all subsequent event-aligned analyses.

Behavioral reaching epochs were concluded within a 400 ms period. We initially extracted wide temporal windows (−1 to +4 s) to avoid edge artifacts. As animals demonstrated more variable behavior after the initial reach-and-hold epoch, we utilized a restricted window of −0.5 to +1 s around reach initiation, encompassing the full reach epoch, for analysis. Spike data were agglomerated over trials, then converted to a 1 kHz time series representation with a Gaussian smoothing kernel (σ=20 ms), then normalized by the number of nonempty trials used.

**Supplemental Figure 1:**
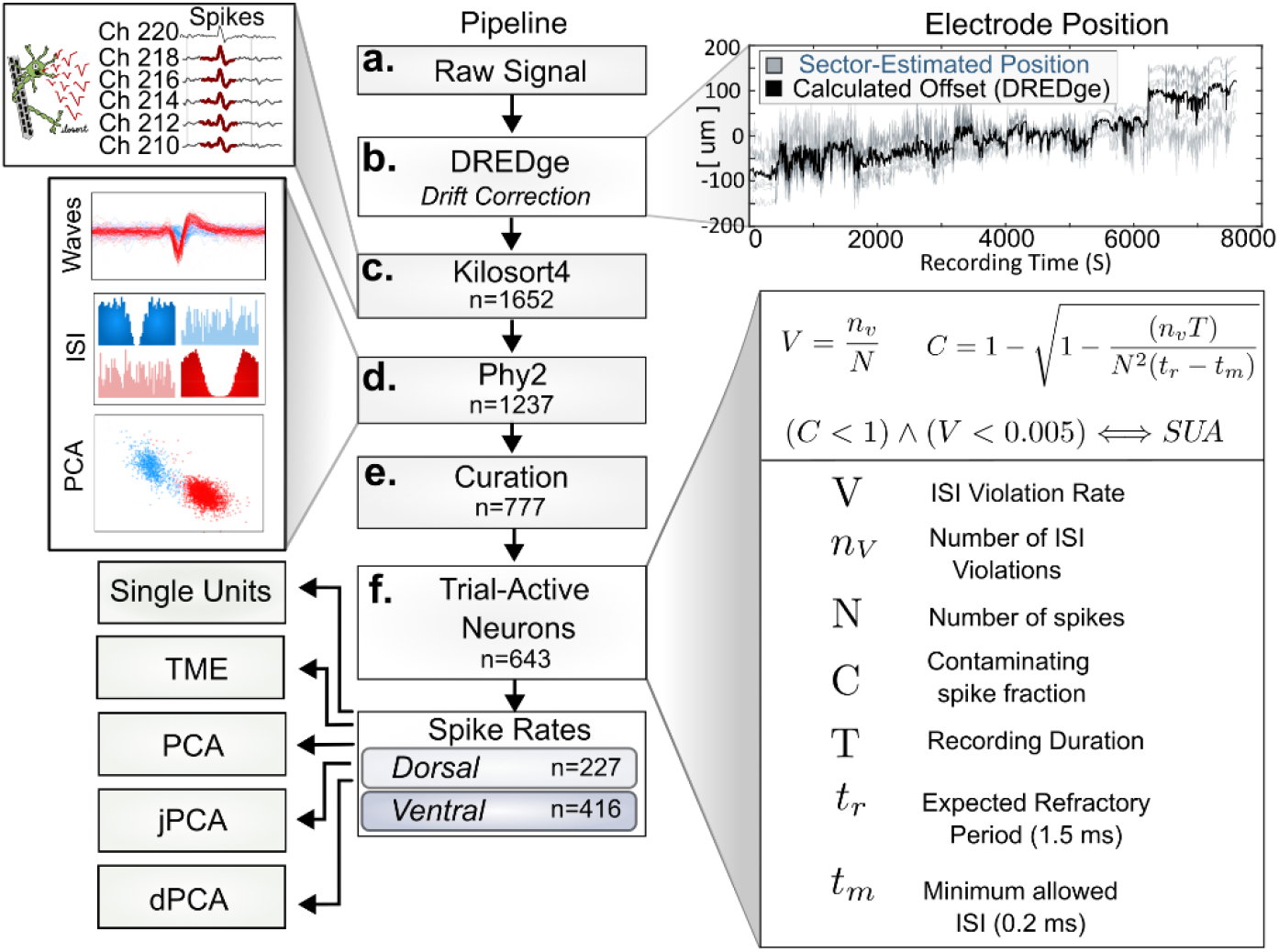
Preprocessing pipeline for processing neural data. A) Raw signals were recorded from the entire length of the probe by a Neuropixels base station via Open-Ephys GUI. B) Raw data was processed through DREDge for linear probe drift correction. C) Spikes were extracted offline by Kilosort4. D) Extracted waveforms were further curated using Phy2 then exported into MATLAB. E) Using custom scripts, we further selected for spikes whose estimated false-positive rate was <5%, as estimated from interspike intervals violations. F) Only trial-active neurons were retained for analysis.

Neural data were sampled across up to 384 channels at 30kHz and recorded using Open Ephys GUI^44^ as a continuous recording as a *.dat* file. Spike data were first drift-corrected using DREDge^10^ and imported into Kilosort4^45^ for automated cluster cutting. Because Kilosort4 has a low false negative rate^45^, we assumed that all meaningful spike waves were extracted. To reduce the instance of false-positives in the dataset, we utilized Phy2 to manually curate recovered spike units. We rejected neurons with odd spike waveforms and units which had high interspike interval violation rates (>5%).

Sorted spikes were imported into MATLAB (MathWorks™) for further curation and analysis (**Supplemental Figure 1**). The percentage of spikes of an assigned neural spike train originating from a contaminating unit (False-Discovery Rate, FDR) was estimated using the non-physiological violations of the observed interspike intervals (ISI)^46^. Units which had an estimated FDR >5%, and had ISI violations >0.5% spikes were excluded for analysis.

#### S2.2 Neural Sampling Depth

**Supplementary Table 1:** Neural sample depth and unit attribution. We utilized data from 8 rats for analysis.

| Rat ID | Reaching Trials (Ipsi) | Reaching Trials (Contra) | Extracted Units (KS4) | Retained Units (Phy2) | Retained Units (Post-hoc) | Trial-Active Neurons | Retained Neurons (Dorsal) | Retained Neurons (Ventral) |
| --- | --- | --- | --- | --- | --- | --- | --- | --- |
| 1J | 20 | 20 | 138 | 101 | 56 | 50 | 12 | 38 |
| 1P | 20 | 20 | 269 | 136 | 64 | 55 | 44 | 11 |
| 2A | 20 | 20 | 297 | 198 | 140 | 127 | 44 | 83 |
| 2G | 20 | 20 | 107 | 122 | 52 | 47 | 25 | 22 |
| 2H | 16 | 16 | 161 | 136 | 107 | 89 | 17 | 72 |
| 2I | 20 | 20 | 263 | 188 | 119 | 73 | 33 | 40 |
| 2K | 34 | 28 | 224 | 129 | 113 | 96 | 34 | 62 |
| 2O | 26 | 34 | 193 | 227 | 146 | 106 | 18 | 88 |
| **TOTAL** | 176 | 178 | 1652 | 1237 | 797 | 643 | 227 | 416 |

Across animals, spike sorting, curation, and filtering yielded a dataset of 797 neurons, with 643 retained for analysis here. Supplemental Table 1 contains the number of neurons recorded and pared from the composite data set by processing step. Notably, ventral neurons were more abundantly sampled than dorsal neurons (416 vs 227), reflecting probe geometry and tissue access. This imbalance in dorsal-ventral sampling was accounted for in downstream analyses via normalization and subsampling where appropriate.

### S3: Single Unit Analysis Properties

To determine whether population-level effects could be explained by single-neuron properties, we quantified firing rate, condition selectivity, and timing at the level of individual neurons.

#### S3.1 Measures of Neural Similarity

We used the trial-averaged PSTH / spike rate vectors for considering the following single-unit neural effects.

1. <u>Spike Rate Waveform Energy (Spike Rate)</u>: This is the root-mean-square (RMS) of the non-normalized PSTH waveform for each neuron. As PSTHs contain the normalized spike rates (x>0), this behaves as an integral of the PSTH waveform, directly correlating to the average number of spikes. The entire PSTH waveform (-500 ms before to 1500 ms after motor onset) was used for calculating RMS.
2. <u>Spike Rate Discriminability (Δ Spike Rate)</u>: Neurons were recorded for two reaching conditions, Ipsilateral and Contralateral. We defined a discrimination index which, for each neuron, corresponds to the difference in PSTH RMS between the reach to the Ipsilateral and contralateral target (**EQ. 1**). Here, bounded [−1 ≤ *x* ≤ +1], a D.I. of +1 corresponds to all waveform energy aligned with the ipsilateral target; a D.I. of -1 corresponds to a complete selectivity in activity for the contralateral target, and a D.I. of 0 corresponds to no difference in average spike activity.

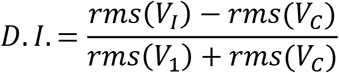

1. <u>PSTH Vector Similarity (Waveform Shape)</u>: Averaged PSTHs correspond to vectors of neural activity. We measured similarity between two vectors as the cosine similarity; bounded [0 ≤ *x* ≤ 1].
2. <u>Peak Spike Activity Timing</u>: From the PSTH, we identified the peak timing of spiking neurons, and evaluated the consistency of this timing by motor condition.

We treated Animal ID, Neural Depth Code, and Reaching Condition as independent variables when evaluating spike effects. Neural PSTHs were pooled, then grouped according to null test hypotheses (e.g., “*H0: Spike-rate distributions do not differ across rats*”). Because sample distributions were not normal, we used non-parametric tests to evaluate differences between groups (**Supplemental Table 2**).

#### S3.2 PSTH Spike-Rate, Discriminability, and Vector Similarity

We considered the discriminability of individual neural PSTHs by reaching condition, dorsal/ventral depth, and rat. We observed that some neurons selectively fired in a single condition. Single unit analysis identified that 145/643 (∼22.6%) of neurons only spiked during a single reaching condition (i.e., had a D.I. score of 1 or -1; were maximally discriminable; **Supplemental Table 2**). To investigate the instability of the results by neurons which only fired in a single condition (and hence had a D.I. value of +-1 or -1), we performed single-neuron analysis twice: 1) with all neurons, 2) with neurons which only fired in both (“Both”) motor conditions.

**Supplemental Table 2:** Count of the number of neurons which only fired in a single motor condition (i.e., had a D.I. of 1 or -1), across rats, relative to the number of total recovered neuron which fired in at least that condition.

| Rat ID | Ipsi (ONLY) | Contra (ONLY) | Neurons (Total) | Ipsi (%) | Contra (%) |
| --- | --- | --- | --- | --- | --- |
| 1J | 1 | 9 | 50 | 2% | 18% |
| 1P | 12 | 14 | 55 | 22% | 25% |
| 2A | 9 | 10 | 127 | 7% | 8% |
| 2G | 2 | 11 | 47 | 4% | 23% |
| 2H | 7 | 14 | 89 | 8% | 16% |
| 2I | 2 | 25 | 73 | 3% | 34% |
| 2K | 5 | 6 | 96 | 5% | 6% |
| 2O | 9 | 9 | 106 | 8% | 8% |
| *Total* | 47 | 98 | 643 | 7% | 15% |

No differences were detected in the spike-rate discriminability (D.I.) across rats or neural depth (**Supplemental Table 3**). These observations were retained when utilizing neurons from the entire data set or only neurons which fired in both motor conditions.

**Supplemental Table 3:**
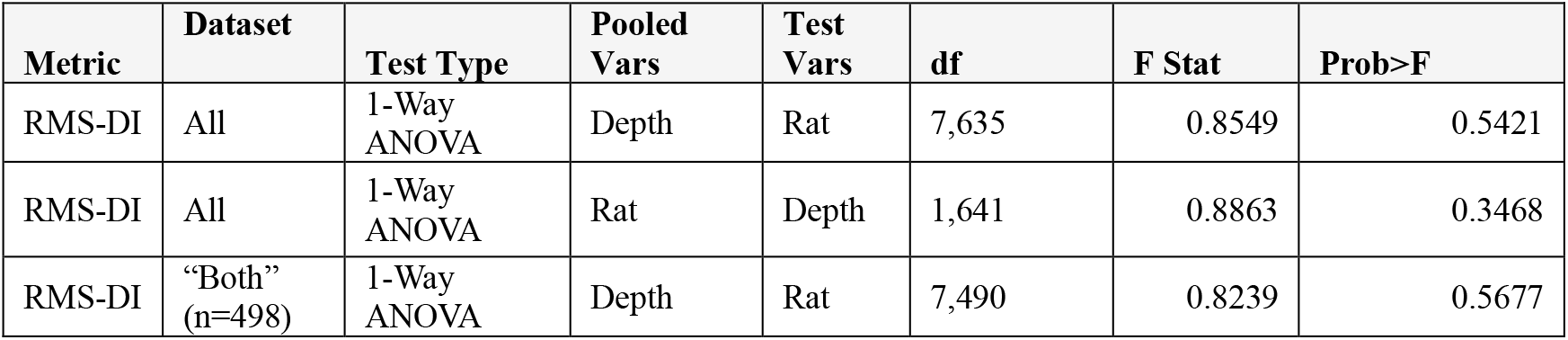
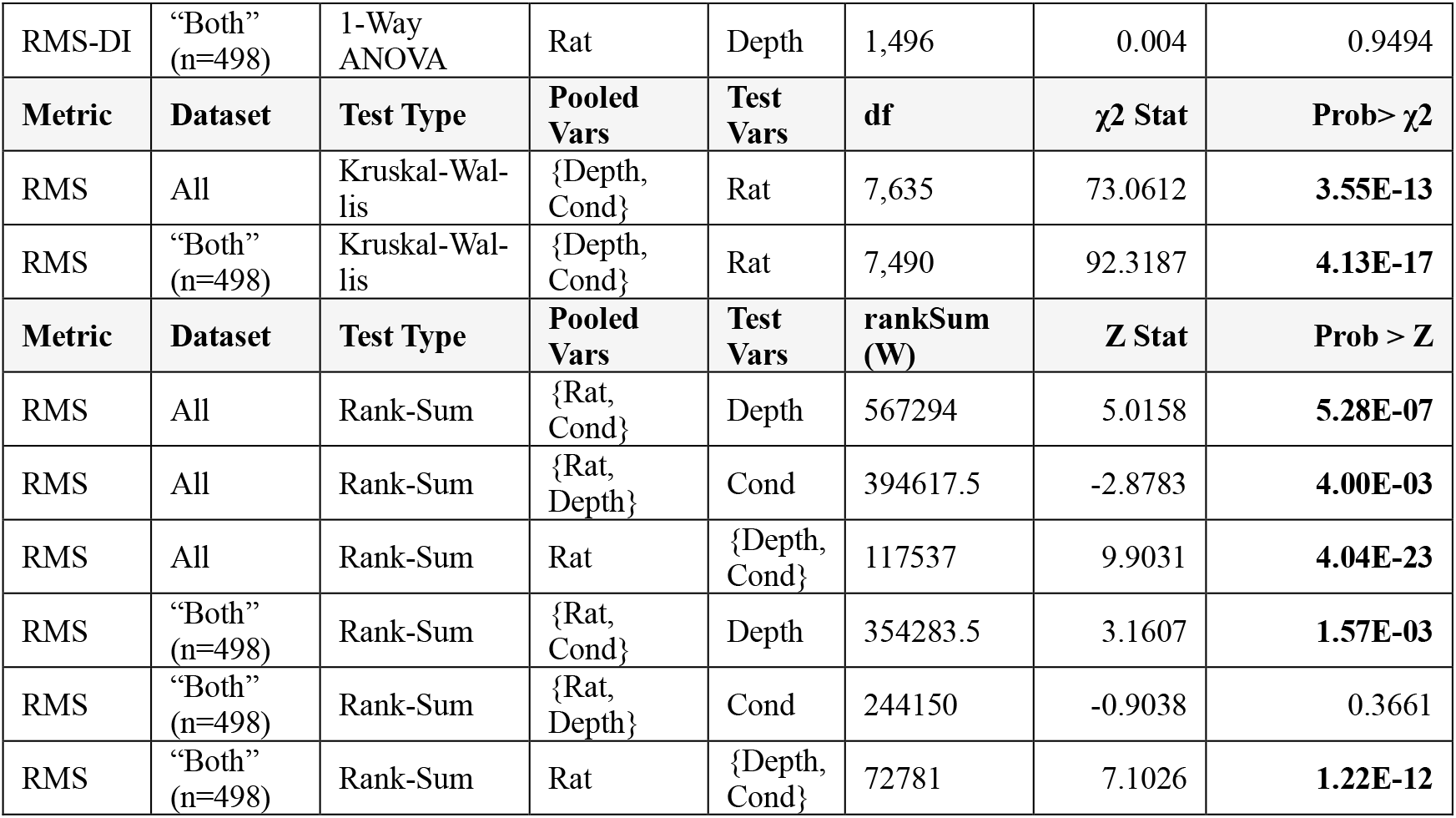
Significance tests of neural spike rates (measured by waveform energy [Root Mean-Square; RMS]) by different groupings and statistical tests. Here, neural ensemble data was grouped according to a test variable (Animal ID [Rat], Depth Code [Depth]), and analyzed using a between-groups test (ANOVA, Kruskal-Wallis, Rank-Sum). Stats were performed for ‘All’ data (n=643 neurons) and neurons which only fired in *both* (‘both’) motor conditions (n=498).

We found that ensemble data were collected across a diversity of neural spike patterns and dynamics ranges; divisions between rat, depth, and motor condition by spike rates RMS all reported as significant (Permutation 1-Way Z-Tests; p=0-0.036). The only exception was when pooling across rats and depths, and testing ensemble differences in motor condition (p=0.3661) by spike rate RMS (**Supplemental Table 3**).

#### S3.3 Spike Peak Timing Analysis

To determine if neural spike timing (tuning) was preserved across ipsilateral and contralateral motor conditions, and whether this varied by dorsal or ventral tag, we treated these peak timings as independent variables and evaluated correlation. Only neurons which spiked in both conditions during the analysis epoch (-500 ms to +1000 ms around movement onset) were used for modeling and plotting.

For visualization, we produced an adjacency matrix of the peak spike timing in the ipsilateral vs. contralateral position (as in **Fig 4**), with a granularity of 1 ms/bin. A 2D smoothing kernel 25ms/σ was used for visualization of the adjacency matrix as an image. We observed that most data were clustered near the matrix diagonal, suggesting similarity in the neural peak timing in both the ipsilateral and contralateral conditions.

To test the concordance of paired peak timing, we divided the 2D adjacency array into 9 regions (Pre-motor, -500 ms to 0 ms; Motor; 0 ms to +250 ms; Post-Motor +250ms to +1000 ms, Ipsilateral/Contralateral lever reach). Neural paired-peak counts were collected for each epoch region for 1) All neurons, 2) dorsal-coded neurons, and 3) ventral-coded neurons.

The distributions of peak-paired timings within these 9 regions differed between dorsal and ventral neurons [χ2 (8, n=2) =18.39, p=0.0185]. The ventral population of neurons had more peaks-pairs within concordant epochs (diagonals) (n =174/342; 50.9%) than the dorsal population (n=69/159; 43.4%).

Having previously observed that the motor active period of both motor conditions usually resolved within 250 ms of onset, we tested the consistency of neurons peak pairs specifically within this motor-active region. Thus, we only considered neurons which spiked during this period (Dorsal: n=151; Ventral: n=336). Contralateral-reach and Ipsilateral-reach peak timings within this interval were correlated using a linear model (**Supplemental Table 4**).

**Supplemental Table 4:** Peak timings of the ipsilateral vs. contralateral neurons within the motor active period (0 to 250 ms after movement onset) were fitted with a linear model. (X = Ipsilateral; Y = Contralateral). Here, the peak timing of neurons in the ipsilateral or contralateral reach is weakly, but significantly conserved. The linear model using dorsal neurons had higher explanatory power for dorsal (22.5%) than ventral (1.3%) neural populations.

| Neural Subset | # Neurons | Slope | Intercept | R2 | RMSE | pSlope | pIntercept |
| --- | --- | --- | --- | --- | --- | --- | --- |
| All | 487 | 0.27 | 74.2 | 0.063 | 67.43 | **0.00367** | **<0.001** |
| Dorsal | 151 | 0.55 | 57.4 | 0.225 | 64.12 | **<0.001** | **0.005** |
| Ventral | 336 | 0.12 | 85 | 0.013 | 68.69 | 0.309 | **<0.001** |

Peak timing was weakly but significantly related across conditions for the ensemble (R²=0.063) and dorsal neurons (R²=0.225), but not for ventral neurons (R²=0.013). The fitted slope term was only significant for the ensemble (p=0.0037) and dorsal-only (p<0.001) datasets, while the y-intercept terms were significant for the ensemble (p<0.001), dorsal-only (p=0.005), and ventral-only (p=0.001) datasets.

### S4: Extended Neuron Dynamics Observations

To test the robustness of populations dynamics across animals, subsamples, and analysis choices, we performed additional tests on different groupings of the data.

#### S4.1 Ensemble Data Demonstrated Rotations across Dorsal and Ventral Populations (TME)

Before we sought to describe potential neural dynamics within the observed low dimensional state, we wanted to ensure that any extracted dynamics ‘features’ were not an artifact of neural preprocessing or chance sampling. Thus, we utilized the Tensor Maximum Entropy (TME)^47^ resampling method on the data tensor of neural spike rate PSTHs. The observed R2 statistic of the neural ensemble data tensor was significantly higher than those generated from 1000 TME (Time, Neuron, Condition; TNC) surrogate values indicating the ensemble (1-tailed t-test, R2=0.6, p=0.002), dorsal (1-tailed t-test, R2=0.51, p=0.003), and ventral (1-tailed t-test, R2=0.59, p=0.002) neural populations could be via a linear dynamical system better than would be expected by chance.

#### S4.2 Effect of Trial-Sampling on jPCA Alignment

To assess how recovered population dynamics depended on specific trial inclusion, for each animal, we drew 1000 bootstrapped surrogates of 10 trials (with replacement) containing trial-wise neural firing rates. As with the ensemble dataset, spike rates within each trial sample were averaged together (i.e., sample averaged spike rates), and used for jPCA.

For each of 1000 shuffles, we also back-projected the sample-trained jPCA matrices into spike rates (i.e., the estimation of what neural spike rates *would* be, given the sample-trained encoding/decoding) from 1) the full set of neurons (per rat), 2) the sampled neurons, 3) the left-out neurons. (“*How well can the sample-trained jPCA matrices encode and decode spike rates across different trials?”)* We observed the VAF of the back projection from sample-trained dataset was effective for encoding/decoding the sample, ensemble, and held-out neurons. We evaluated the canonical correlations (**Sup. Fig 2A**), linear axis correlation (**Sup. Fig. 2B-C**), and variance-accounted (**Sup Fig 2D**) for by sample group.

Next, we determined the trial-by-trial jPCA latent stability. We used each rat’s ensemble-trained jPCA matrices to project expected jPCA latents using single-trial spike rates (**Supplemental Fig 2E**). Even without alignment, most rats maintained distinguishable latent drives which were across the trial samples, indicating a gross conservation in the patterns of the neural activity comprising these drives across trials.

**Supplemental Figure 2:**
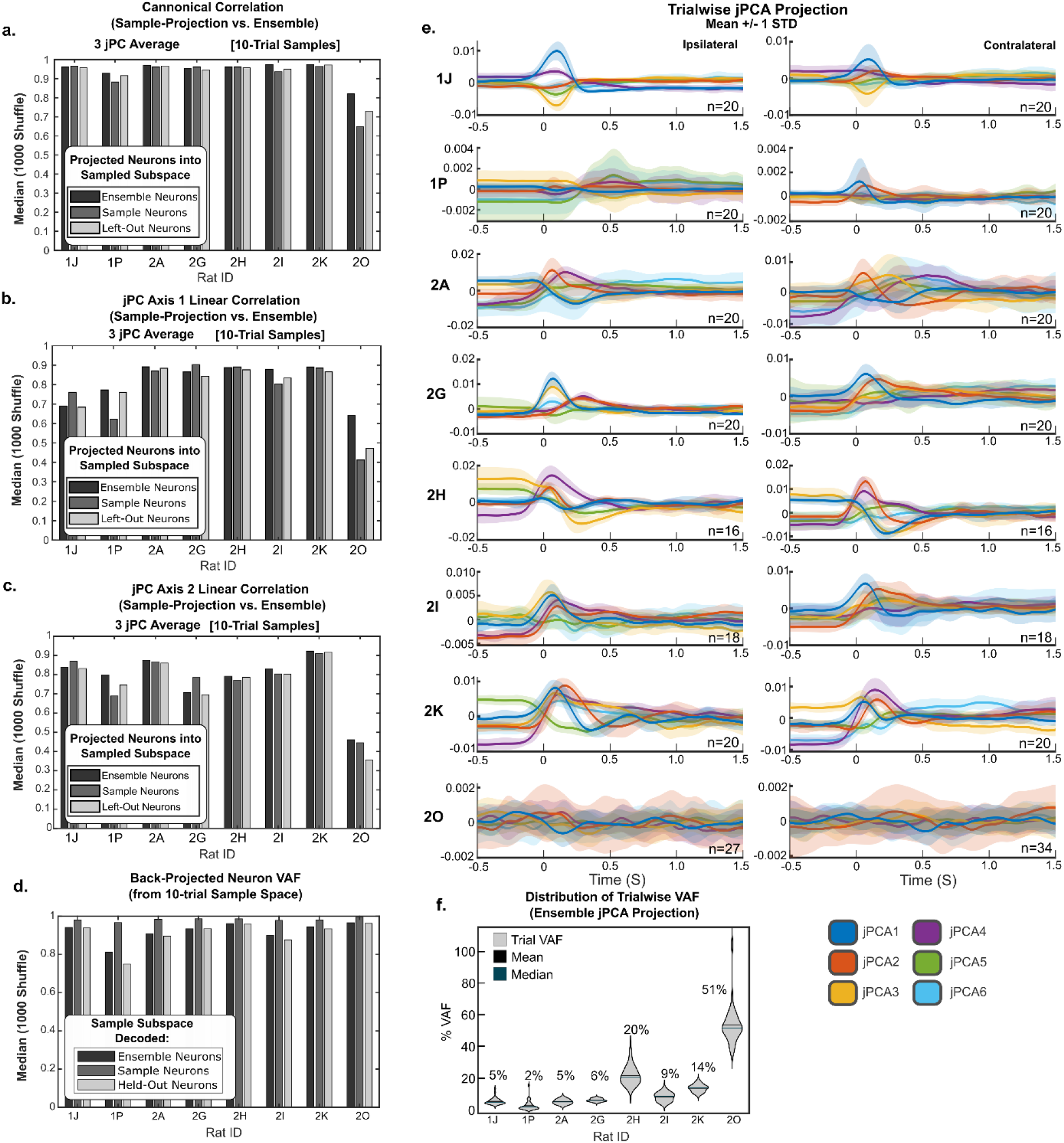
Effect of Trial Selection on jPCA Alignment. For each rat, 1000 surrogate draws of 10 trials were used producing averaged spike rate and subjected to jPCA. jPCA subspaces of each surrogate were aligned against the jPCA subspace generated by the ensemble of all trials for the rat. The distance between each surrogate to the ensemble was measured via **A**) canonical correlation and **B-C**) jPC axis linear correlation metrics. D) Sample-trained jPCA matrices were used to first project jPCA latents from ensemble, sample, and held-out neural spike rates, then back-project expected spike rates from those latents (encoding/decoding). Across the median of the shuffles, we observed a high VAF of reconstructed spike rates within the 6-dimensional jPC space within the 10-trial samples regardless of whether spike data emerged from the average of the ensemble, sample, or held-out data. **E**) Overlay of the average ± 1 standard deviation of the projection of the trial-ensemble trained jPCA (on a per-rat basis) for each trial. Here, most rats maintained consistent neural recruitments across trials, indicated by similar jPCA series projections across the ensemble. **F**) Trialwise variance-accounted-for each trial (no sample-averaging) utilizing the jPCA structures trained on the per-rat ensemble. Here, VAF was lower than for the sample-averaging, consistent with the relative sparsity of neural responses on a single-trial basis. Rat 2O forms and outlier, having relatively flat neural dynamics, without much trialwise variation, thus capturing much of this ‘flat’ activity.

#### S4.3 Effect of Rat-Sampling on jPCA Alignment

We compared the jPCA analyzes generated from a single-rat basis by both comparing individual rats to the ensemble and jackknifing each rat’s neurons from the ensemble. Similarity was measured as canonical correlation relative to the ensemble (**Supplemental Table 5**).

**Supplemental Table 5:** Groupwise canonical correlation of rat jPCA to the ensemble jPCA (above). Canonical correlation of the jPCA *without* a rat to the jPCA of the ensemble (rat jackknife).

| Rat ID | Per-rat canonical correlation($\rho$) | | | | | | | |
| --- | --- | --- | --- | --- | --- | --- | --- | --- |
|  | 1J | 1P | 2A | 2G | 2H | 2I | 2K | 2O |
| Plane |  |  |  |  |  |  |  |  |
| 1 | 0.871 | 0.652 | 0.920 | 0.858 | 0.932 | 0.803 | 0.784 | 0.530 |
| 2 | 0.746 | 0.437 | 0.888 | 0.679 | 0.689 | 0.881 | 0.710 | 0.561 |
| 3 | 0.620 | 0.963 | 0.964 | 0.945 | 0.779 | 0.952 | 0.873 | 0.565 |
| Rat ID | Without-rat canonical correlation($\rho$) | | | | | | | |
|  | ~1J | ~1P | ~2A | ~2G | ~2H | ~2I | ~2K | ~2O |
| Plane |  |  |  |  |  |  |  |  |
| 1 | 0.999 | 1.000 | 0.974 | 0.640 | 0.923 | 0.984 | 0.878 | 1.000 |
| 2 | 0.797 | 1.000 | 0.970 | 0.642 | 0.891 | 0.855 | 0.990 | 1.000 |
| 3 | 0.740 | 1.000 | 0.983 | 0.999 | 0.943 | 0.700 | 0.998 | 1.000 |

#### S4.4 Effect of Depth-Sampling on jPCA Alignment

To test whether the observed alignment exceeded that expected by chance, we compared observed dorsal (n=227) and ventral (n=416) jPCA subspaces to sample-size-matched surrogate populations drawn from the full dataset. As with the trial-sampling, we compared the similarity of the extracted jPCA dynamical motifs using 1) canonical correlation and 2) axis correlation after Procrustes alignment. To evaluate significance of these depth-isolated alignments relative to chance across the ensemble, we generated 5000 sample-sized controlled surrogate data sets (i.e., draw without replacement) of the spike rate data, performed jPCA, and measured alignment to the jPCA surrogates to the *ensemble* dataset. We then measured canonical and linear axis correlation via a two-tailed deviation of the observed test statistic vs surrogate population.

As with canonical correlation, aligned linear correlation was high across all 6 axes for both the dorsal (0.79-0.97) and ventral (0.98-0.99) groups relative to the ensemble structure (**Supplemental Table 6**). Ventral neurons were often more highly correlated with the ensemble than could be created with any admixture of the dorsal and ventral neurons (surrogates), (p= 0-0.014), supporting a finding that the larger ventral pool of neurons define the extracted jPCA dynamics across the *ensemble*, but that these dynamics are shared in the dorsal pool. Across all metrics, dorsal and ventral subspaces closely matched the ensemble structure, with alignment exceeding that of surrogate populations.

**Supplemental Table 6:** Linear axis correlations between jPCA subspaces on the ensemble to the Procrustes-aligned subspaces for a defined neural subgrouping (dorsal, ventral). Significance was determined relative to a sample-size balanced set of 5000 surrogate statistics, randomly drawn from the ensemble. Ventral neurons had a significantly higher axis correlation than would be expected from a number-balanced sample. In contrast, the dorsal neurons did not significantly differ, indicating that the dorsal set was no better or worse of a fit than the average admixture of dorsal and ventral neurons.

| Group-<br>ing | Plane | Test Val | 95%<br>Shuff CI<br>(Lower) | 95%<br>Shuff CI<br>(Upper) | pVal |
| --- | --- | --- | --- | --- | --- |
| Ventral | JP1 [1] | 0.997 | 0.137 | 0.981 | **0** |
|  | JP1 [2] | 0.998 | 0.281 | 0.915 | **0** |
|  | JP2 [3] | 0.991 | -0.555 | 0.978 | **0.004** |
|  | JP2 [4] | 0.993 | 0.419 | 0.985 | **0.014** |
|  | JP3 [5] | 0.991 | 0.695 | 0.980 | **0.006** |
|  | JP3 [6] | 0.990 | -0.024 | 0.964 | **0.0064** |
| Dorsal | JP1 [1] | 0.851 | 0.116 | 0.985 | 0.7904 |
|  | JP1 [2] | 0.832 | 0.243 | 0.976 | 0.836 |
|  | JP2 [3] | 0.790 | -0.627 | 0.981 | 0.7812 |
|  | JP2 [4] | 0.973 | 0.487 | 0.990 | 0.2464 |
|  | JP3 [5] | 0.868 | 0.608 | 0.988 | 0.6508 |
|  | JP3 [6] | 0.957 | -0.707 | 0.984 | 0.2236 |

#### S4.5 Effect of Mean-Subtraction on dPCA Alignment

Mean subtraction of neural spike traces provides a per-channel offset, centering the data, and evaluating changes relative to this offset. However, we determined that dPCA was effective even without this biasing factor. In practice, the mean-subtraction had minimal impact on the *time* marginalizations, (save that the components were now centered around 0, rather than starting at 0), and distributed variance between the *condition* and *time×condition* interaction marginals. The *time* marginalization dominated across the neural ensemble, accounting for 57.3% of the variance, with *condition* (∼6–7 VAF%) and *time×condition* interactions (∼3 VAF%) explaining comparatively little.

As with the mean-subtracted case, the canonical correlation between dorsal/ventral dPCA and the ensemble were high (all 0.9953-1.0). Because we were no longer performing a per-channel, per condition offset, performing dPCA separately for the dorsal and ventral populations resulted in a higher variance explained by ipsilateral/contralateral reaching *condition* (D: 22.1%, V: 19.8%) whereas this was previously trivial.

We found that the intersections within each group were highly conserved across marginals (All: Jaccard=0.45–0.64; Dorsal: 0.66–0.85; Ventral: 0.51–0.71; all p<10⁻⁶, FDR-corrected), and neuron-wise contribution magnitudes were strongly correlated (Spearman ρ=0.64–0.88). Similarly, there was no significant overlap enrichment between dorsal and ventral groups in any marginalization (Jaccard=0.13–0.22; FDR-corrected p>0.17).

Even when geometrically aligning subspaces via Procrustes alignment and measuring axes correlation, dorsal and ventral subspaces remained substantially similar to the ensemble. Ventral neurons were often the best-fit tested group (p=0.000) than any admixture of dorsal and ventral neuron (i.e., surrogate), indicating that these neurons best fit the dynamics patterns extracted across the ensemble.

Similarly, the canonical correlation of the *time* dPCA marginals evaluated on a per-rat basis relative to the ensemble was also high without mean-subtraction (**Supplemental Table 7**). Rat 2O was the most divergent (ρ>0.91), but nonetheless showed a high fit with dPCA.

**Supplemental Table 7:** Canonical correlation alignment statistics of animal-wise subspaces aligned with the ensemble without neuron-wise mean subtraction. Canonical correlations are high, indicating near linearity of animal-wise and ensemble dPCA subspaces, even with fewer neurons.

| dPCA Marginal | 1J | 1P | 2A | 2G | 2H | 2I | 2K | 2O |
| --- | --- | --- | --- | --- | --- | --- | --- | --- |
| *time* | 0.996 | 0.994 | 1.000 | 0.999 | 1.000 | 0.999 | 1.000 | 0.952 |
| *cond* | 0.992 | 0.995 | 0.997 | 0.993 | 0.999 | 0.994 | 0.997 | 0.998 |
| *timex-cond* | 0.989 | 0.968 | 0.990 | 0.995 | 0.993 | 0.974 | 0.984 | 0.919 |

Axis-wise correlations and Procrustes residuals likewise showed minimal deviation across animals. Thus, dorsal and ventral subsamples of the data could be well-described by each-other’s subspace and the effect of mean subtraction did not substantially vary results.

#### S4.6 Discrete-Time-Warped (DTW) Alignment of dPCA Marginals

We noted that some components, such as dPC-T3 may be better fit across depths with a change in temporal progression, rather than a lag. To determine whether dorsal-ventral differences were better described by changes in temporal progression rather than a fixed delay, components were additionally aligned using dynamic time warping (DTW). While most components were adequately described by a fixed lag, dPC-T3 (**Fig. 6D**) achieved substantially higher alignment following DTW (R=0.85 vs R=0.76, p=0.001 Ipsi; R=0.85 vs. R=0.72, p=0.342, Contra) compared to lag alignment. Confidence intervals were nevertheless broad, indicating the best fitting of this component may vary by included trials.

